# A collection of patient-derived intestinal organoid lines reveals epithelial phenotypes associated with genetic drivers of pediatric inflammatory bowel disease

**DOI:** 10.1101/2025.06.11.659052

**Authors:** Zahra Shojaei Jeshvaghani, Carmen Argmann, Michal Mokry, Johan H. van Es, Maaike H. de Vries, Lauren V. Collen, Daniel Kotlarz, Mia Sveen, Phillip Comella, Scott B. Snapper, Christoph Klein, Eric Schadt, Hans Clevers, Ewart Kuijk, Edward Nieuwenhuis

**Author notes:** contributed equally (Ewart Kuijk, Edward Nieuwenhuis).

## Abstract

Pediatric Inflammatory Bowel Disease (IBD) is a chronic condition characterized by persistent intestinal inflammation in children and adolescents. Despite a rising global incidence, the underlying causes and optimal management strategies for pediatric IBD are still not fully understood. Compared to adult IBD, pediatric IBD frequently presents with distinct disease phenotypes, and is more commonly linked to rare monogenic variants that cause intestinal epithelial barrier dysfunction or affect the function of mucosal immune cells. While more than 100 genes have been associated with early-onset IBD, the roles of many of these genes in the intestinal epithelium and the mechanisms by which genetic variants contribute to disease remain poorly defined.

Here we aimed to improve our understanding of intestinal epithelium dysfunction in early-onset IBD by conducting extensive molecular and cellular characterization to gain insights into patient-specific epithelial phenotypes and identify therapeutic targets. We generated intestinal epithelial organoids (IEOs) from 94 pediatric IBD patients, representing diverse clinical characteristics and including those with monogenic variants (*BTK* n=4, *TTC7A* n=3, *IL10RA* n=1, *LRBA* n=1, *STXBP2* n=1, *TTC37* n=1, *TRNT1* n=1, *PLCG2* n=1, *DKC1* n=1, *POLA1* n=1), and 46 non-IBD controls. This effort resulted in the largest RNA-seq dataset of pediatric IBD intestinal epithelial organoids to date, encompassing both baseline conditions and post-immunological stimulation, serving as a valuable resource for future research.

We observed that IEOs effectively initiate inflammation upon stimulation with bacterial lysate, regardless of disease status, origin, or mutation status. Inflammatory stimulation triggered single-gene upregulation of IBD-linked *SERPINA1* and *LIFR* across the IBD population compared to controls, suggesting their role in intestinal epithelial innate immune responses. However, co-expression network analysis showed no consistent transcriptional signatures across the entire IBD group at the systems level. Instead, differences emerged between controls and specific genotypes (*TTC7A*, *STXBP2*, *LRBA*), with *STXBP2* and *LRBA* sharing an upregulated transcriptional response of IL-1 and SLC30-mediated zinc trafficking pathways.

These findings underscore the potential of IEOs as a valuable model for studying IBD and offer key insights that could guide the development of targeted therapies for both monogenic and non-monogenic forms of IBD.

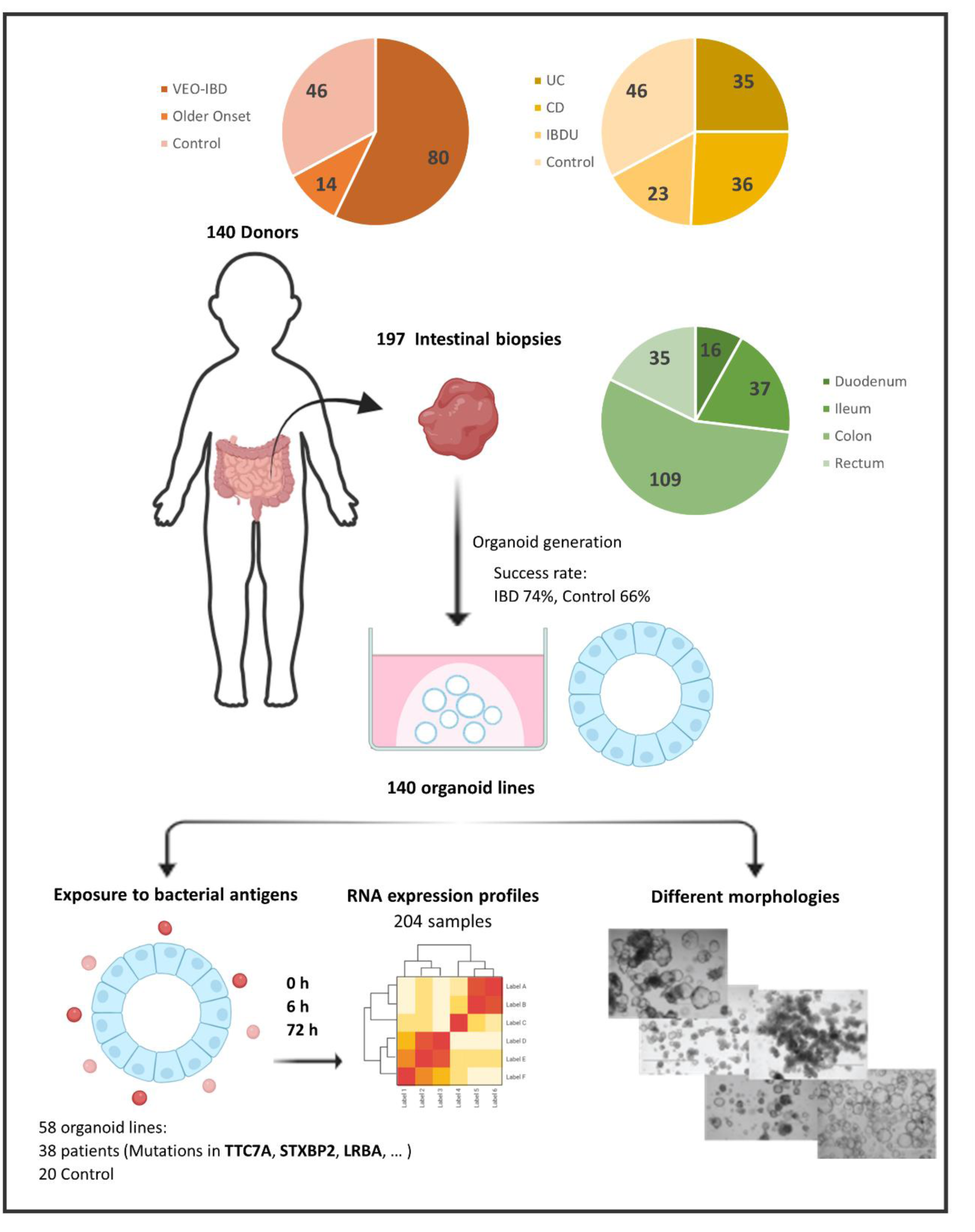

## Introduction

Inflammatory bowel disease (IBD) is characterized by chronic inflammation of the gastrointestinal tract. Based on the histopathological characteristics, IBD can be categorized as Crohn’s disease (CD), ulcerative colitis (UC), and IBD-unclassified (IBDU).^1^ In pediatric cases, IBD typically manifests before the age of 17, per the Paris classification.^2^ The incidence and prevalence of IBD is increasing globally among both children and adults.^3–5^ Early-onset IBD is highly heterogeneous, driven by a complex interplay of aberrant intestinal immune responses, epithelial barrier dysfunction, gut microbes, and environmental factors in genetically predisposed individuals.^6,7^ Despite the availability of numerous IBD treatments, effective therapies remain elusive for many patients with very early-onset IBD (VEO-IBD). This is largely because most therapies were validated in large cohort studies of adult IBD, making their effectiveness in individual cases, particularly those with VEO-IBD with monogenic causes, unpredictable. This issue is further complicated by the inherent differences between VEO-IBD and IBD diagnosed later in life. VEO-IBD presents unique challenges due to its low prevalence and largely unknown etiology.

While VEO-IBD patients are thought to have greater genetic contribution to disease development, definitive monogenic diagnoses are identified in fewer than 10% of cases based on large cohorts with universal access to whole exome sequencing.^8,9^

A key player in IBD is the intestinal epithelium, i.e., the monolayer of epithelial cells that form the luminal surface of the gastrointestinal tract.^10,11^ Inflammation is initiated when epithelial integrity is lost, allowing microbiological molecules to pass the epithelial barrier thereby triggering an immune response that is not adequately resolved in IBD patients. Over 100 genes have been linked to monogenic forms of IBD, with most genes associated with immune disorders.^12^ However, mutations in genes like, *TTC7A*, *STXBP2*, *STXBP3*, *COL7A1, FERMT1, ITGA6, ITGB4*, *SLC9A3*, *GUCY2C*, *SLC26A3*, *EPCAM*, *IKBKG*, *TTC37*, *CD55*, *ADAM17*, and *SKIV2L* cause IBD by interfering with intestinal epithelium homeostasis, underscoring the role of the epithelium in disease development.^13^

Intestinal epithelial organoids (IEOs) are three-dimensional (3D) structures derived from intestinal stem cells closely resembling the intestinal epithelium. Unlike animal models and immortalized cell lines, which poorly replicate the human intestinal niche, 3D intestinal organoids offer a more accurate platform for studying primary epithelial cells under near-physiological conditions.^14,15^ This unique model, with its exposed basolateral side, is ideal for studying IBD disease pathology, where the loss of intestinal barrier integrity leads to the basolateral exposure of epithelial cells to bacterial antigens, triggering the inflammatory responses characteristic of IBD. IEOs have been widely used in IBD research to study host-microbe interactions, barrier function, epithelial damage, drug discovery, personalized medicine, and the genetic defects underlying IBD pathogenesis.^16–22^ While organoids have been shown to maintain genomic stability and key disease characteristics,^23–26^ studies on colon organoids derived from UC patients indicate a gradual loss of the inflammatory profile observed in fresh biopsies, with inflammation becoming undetectable after seven days.^27,28^ This shift to an uninflamed state in isolated cultures underscores the need to effectively reestablish inflammation, a hallmark of IBD, to create a representative model before exploring other underlying mechanisms. However, this normalization also provides an opportunity to study dysregulated immune responses to microbial stimulation, a key aspect of IBD pathogenesis, particularly those mediated by the intestinal epithelium, independent of the initial inflammatory state of the biopsy.^7,29^

Here, we aim to better understand intestinal epithelium dysfunction in pediatric IBD using organoid technology, and to identify patient-specific epithelial phenotypes and potential intervention strategies. As part of an ongoing effort, we generated IEOs from a cohort of 197 biopsies collected from various anatomical intestinal locations and inflammatory states, derived from control and pediatric IBD patients with diverse clinical profiles. Here, we present the first results of this study, where we performed RNA sequencing on 38 IBD and 20 control organoid lines under unexposed conditions and when exposed to bacterial antigens, demonstrating that IEOs effectively model inflammation when exposed to bacterial antigens. This analysis identified distinct single-gene epithelial phenotypes in IBD organoids compared to controls following inflammatory stimulation, along with gene expression subnetworks associated with bacterial stimulation and specific monogenic variants. These findings offer valuable insights that could inform the development of genotype-specific targeted therapies.

## Materials and Methods

### Research approval

Biopsy samples from clinically required surgical resections or diagnostic endoscopy, were collected from the patients and control subjects, along with clinical information, with their prior informed consent. Data collection and research studies were conducted in adherence to the ethical guidelines established by the ethics committees. 160 Biopsies were acquired from Boston Children Hospital (protocol # IRB-P00000529, titled: Pediatric Gastrointestinal Disease Biospecimen Repository and Data Registry), 32 from Klinikum der Universitat Munchen (KUM protocol # 806-16, titled: Analysis of immunological and genetic causes in pediatric inflammatory bowel disease) and 5 biopsies from the University Medical Center Utrecht (Medisch Ethische Toetsings Commissie (METC) protocol # 10/402, titled: Specific Tissue Engineering in Medicine (STEM)).

### Clinical data

Clinical data were obtained from anonymized medical records of the study population. Donors were classified into three groups: VEO-IBD, older-onset IBD, and controls, based on disease status and age at diagnosis. The control group included healthy donors or individuals diagnosed with non-IBD gastrointestinal conditions. For both IBD and control groups, the following variables were collected: age at sample collection, gender, sample origin, and inflammation status. Additional data were recorded for the IBD group, including age at diagnosis, IBD subtype (CD, UC, or IBDU), mutations in known IBD-associated genes, macroscopic disease location, perianal involvement, stricturing status in CD and IBDU patients, extraintestinal manifestations, history of IBD-related surgery, and medication history. A detailed summary is provided in **Table 1** and **Supplementary Tables 1 and 2**.

**TABLE 1.** Patient and Biopsies Characteristics.

|  | VEO | Older Onset | Control |
| --- | --- | --- | --- |
| **Patients, n** | **80** | **14** | **46** |
| Male/female ratio | 1.35 | 1.33 | 1.42 |
| Age at diagnosis, mean (±SD), years | 2.9 (1.8) | 9.5 (3.6) | NA |
| Age at sample collection, mean (±SD), years | 8 (5.8) | 11.8 (4.4) | 6.9 (7.5) |
| **Diagnosis, n (%)** |  |  |  |
| UC | 34 (42.5%) | 1 (7%) | NA |
| CD | 26 (32.5%) | 10 (71.5%) | NA |
| IBDU | 20 (25%) | 3 (21.5%) | NA |
| **Disease Location, n (%)** |  |  |  |
| Colonic only | 49 (68%) | 5 (45.5%) | NA |
| Ileocolonic | 9 (12.5%) | 5 (45.5%) | NA |
| Panenteric<sup>a</sup> | 14 (19.5%) | 1 (9%) | NA |
| **Surgical History, n (%)** |  |  |  |
| Bowel resections | 5 (8%) | 2 (17%) | NA |
| Ostomy | 4 (6%) | 1 (8%) | NA |
| Both | 6 (9%) | 0 (0%) | NA |
| **Strictureing<sup>b</sup>, n (%)** | 10 (27%) | 3 (30%) | NA |
| **Perianal Involvement, n (%)** | 18 (24%) | 3 (23%) | NA |
| **Monogenic IBD Mutation, n (%)** | 8 (10%) | 7 (50%) | NA |
| **Extraintestinal Comorbidities, n (%)** | 23 (34%) | 6 (50%) | NA |
| **Total No. Biopsies, n (%)** | **111** | **22** | **64** |
| **Sample location of origin, n (%)** |  |  |  |
| Duodenum | 9 (8%) | 1 (5%) | 6 (10%) |
| Ileum | 20 (18%) | 4 (18%) | 13 (20%) |
| Colon | 62 (56%) | 13 (59%) | 34 (53%) |
| Rectum | 20 (18%) | 4 (18%) | 11 (17%) |

<sup>a</sup> Panenteric phenotype affects the entire gastrointestinal tract
<sup>b</sup> Not applicable to patients with UC

### Organoid establishment

Biopsies were obtained by flexible gastroduodenoscopy, colonoscopy or resection material from intestinal surgery, collected in cold DMEM supplemented with 10% FBS and Penicillin-Streptomycin, and cryopreserved at –80°C in 90% FBS/10% DMSO (SIGMA, 02650-100ML) within one hour of collection, prior to organoid generation. The isolation of intestinal crypts, as well as the establishment, culture, and cryopreservation/thawing of human intestinal organoid lines from biopsies were performed as previously described.^23,30^ In short, biopsies were washed with cold complete chelation solution and incubated in chelation solution supplemented with 10 mM EDTA for 30 - 60 minutes at 4°C on a rocking platform. The supernatant was harvested, and EDTA was washed away. Crypts were isolated by centrifugation and plated in prewarmed 24-well plates, embedded in droplets of 50-70% Matrigel (growth factor reduced, phenol free; BD bioscience) diluted in basal medium (BM). The BM medium consisted of advanced DMEM/F-12 (Gibco, 12-634-010), Penicillin-Streptomycin 50 U/mL (Gibco, 15070063), 10mM HEPES (Gibco, 15630080) and 1% GlutaMax (Gibco, 15630-056). Matrigel droplets were polymerized for 10 minutes at 37 °C and then immersed in organoid establishment medium composed of BM enriched with 0.5 nM Wnt-surrogate (U-Protein Express, N001-0.5mg), 2% Noggin conditioned medium (NCM) (U-Protein Express, N002 - 100 mL), 20% R-Spondin conditioned medium (derived from RSPO1-expressing HEK293T cells, kindly provided by Dr. C. J. Kuo, Department of Medicine, Stanford, CA), 50 ng/mL human EGF (Peprotech, 315-09-1MG), 10 mM Nicotinamide (SIGMA, N0636-500G), 1.25 mM N-acetyl cysteine (SIGMA, A9165), 1x B27 (Thermo Fisher, 12587010), 500 nM TGF-b inhibitor (A83-01,Tocris Bioscience, 2939/10), 10 μM P38 inhibitor (SB2021190, SIGMA, S7067-25mg), Primocin 100 μg/ml (InvivoGen, ant-pm-1). Once the organoid cultures were established, the organoid establishment medium was replaced with expansion medium (EM), which is identical except for the substitution of the Wnt surrogate and commercial NCM with 50% Wnt3a-conditioned medium and 10% NCM, respectively, both derived from inhouse cell lines. The medium was refreshed every 2-3 days and organoids were passaged every 7-14 days, either by mechanical shearing at a ratio of 1:1 to 1:4, depending on the culture density, or by dissociating them into single cells using TrypLE™ (Thermo Fisher Scientific, 12604021). Single-cell cultures were treated with EM supplemented with 10 µM ROCK inhibitor (Y-27632, Abcam, 120129) for the first 2-3 days. All cultures were kept in incubators at 37°C, in a humidified atmosphere with 5% C0_2_. Morphology and overall condition of organoids was monitored using standard microscopy and the EVOS cell imaging system (Thermo Fisher Scientific).

### Exposure to bacterial antigens

Bacterial lysate was prepared from *Escherichia coli* HST-08 Stellar competent cells. Bacteria were inactivated at 70°C for 1 hour, followed by exposure to-80°C for 2-3 hours. Subsequently, samples were sonicated (Covaris ultrasonicator, Duty cycle: 20%; Intensity: 10; Cycles burst: 500; time: 30 sec), centrifuged at 10.000g, placed at 4°C for 30 minutes followed by filter-sterilization. The samples were incubated at 70°C for an additional 15 minutes to ensure proper heat-inactivation. The amount of bacterial lysate used for organoid exposure was titrated (ranging from 1µL to 20µL lysate per 500µL medium) to determine the concentration that elicits half of the maximum CXCL-8 response at the mRNA level. Seven days post-splitting, both patient-derived and control organoids were exposed to the optimal bacterial lysate concentration for 0, 6, or 72 hours. After exposure, the organoids were immediately lysed in TRIzol® LS (Ambion, 10296-028) and stored at - 80°C until further RNA isolation and library preparation for RNA sequencing.

### RNA sequencing and analysis

Total RNA was isolated with TRIzol® LS (Ambion, 10296-028), according to the manufacturer’s instructions. Polyadenylated mRNA was isolated using Poly(A) Beads (NEXTflex) and sequencing libraries were made using the Rapid Directional RNA-seq kit (NEXTflex). RNA-seq was conducted on 204 samples from 58 distinct organoid lines, derived from 30 unique pediatric IBD donors and 17 unique non-IBD donors. Organoids were sequenced at baseline and after 6 and 72 hours of bacterial stimulation. In some cases, duplicate samples from the same patient were included, and/or multiple intestinal regions (e.g., colon and ileum) were analyzed, resulting in 204 samples **(Supplementary Tables 1 and 2)**.

Sequencing was performed using the Nextseq500 platform (Illumina), producing single-end reads of 75bp and producing 10-15M reads per sample. Reads were aligned to the human reference genome (GRCh37) using STAR.^31^

Differentially expressed genes (DEGs) in stimulated vs. baseline and IBD vs. control organoids were identified using DESeq2 with an adjusted p-value threshold of <0.01 and an absolute log2foldchange threshold of >1.^32^ Functional enrichment was performed on log2foldchange values with the R-package clusterProfiler, using minGSSize = 3 and maxGSSize = 800 and a pvalueCutoff = 0.05 of Bonferroni corrected p-values.^33–35^ For gene set enrichment analysis, the packages gage and gageData were used.^36^

For Weighted gene co-expression network analysis (WGCNA) raw count data was prefiltered to keep genes with CPM>1.0 for at least 30% of the samples (n=13615 genes). After filtering, count data was normalized via the weighted trimmed mean of M-values.^37^ Gene expression matrices were generated using the voom transformation and adjusted for technical variables (e.g. batch and organoid intestinal location) using the *limma* framework **(Supplementary Table 3 and Supplementary Figure 1).**^38^ WGCNA was performed in R on adjusted, log2 counts per million (lcpm)-transformed count data from baseline, 6-hour, and 72-hour time points following bacterial stimulation (n=192 samples, 13615 genes), as previously described.^39,40^ A signed network (β = 12) was generated using blockwiseModules function (maxBlockSize = 30000) with default parameters **(Supplementary Table 4)**. See **Supplementary Figure 2** for scale free plots and clustering analysis. Pathway and gene enrichment analyses for the WGCNA modules were conducted using Fisher’s exact test in R, with Benjamini-Hochberg correction for multiple testing. BioPlanet pathways (sourced from Enrichr, 2022) and (VEO-)IBD-related genes were used as reference sets for pathway and gene enrichment, respectively.^41^

To identify DEGs in samples with a known genotype relative to controls (excluding control donors 8, 13, 26, and 91), we considered bacterial exposure time as an ordered factor. Specifically, we used a nested linear null model: gene_expression ∼ bacterial_exposure_time + batch + location. The model was fitted using lmFit (from the limma R package, version 3.60.6) in R (version 4.4.1).^42^ We then applied eBayes (also from limma) to moderate the standard errors and used topTable to extract statistical significance (including FDR). Genes were considered differentially expressed by ranking on a p-value of <0.05, unless stated otherwise. Technical replicates were limited to N=1, except for D039_BTK and each TTC7A donor, which had N=2 at each time point **(Supplementary Table 1)**.

### Statistical analysis

Statistical analyses of clinical data and organoid growth were conducted using GraphPad QuickCalcs (https://www.graphpad.com/quickcalcs/).

## Results

### Clinical Characteristics

To investigate intestinal epithelial phenotypes, we collected intestinal biopsies and associated clinical data from a cohort of IBD and control donors. For a clear understanding of the study population, **Table 1** details the demographic characteristics of the VEO-IBD, older-onset pediatric IBD, and control groups. Patients with VEO-IBD were diagnosed at younger than 6 years of age (range: 0-5.9 years), and age at time of sample collection from 0.16 to 23 years. Patients with older onset pediatric IBD were diagnosed between age 6 years and 18 years, and age at time of sample collection ranged from 6 to 19 years. To explore the age-specific variability in IBD clinical characteristics, we compared the overall pediatric IBD population (VEO-IBD vs. older-onset). Our findings reveal significant associations between the age of disease onset (VEO-IBD vs. Older Onset) and diagnosis (UC, CD, IBDU), as well as disease location and genetic factors. Notably, the majority of VEO-IBD patients (∼43%) were diagnosed with UC, with disease confined to the colon. In contrast, older-onset patients (∼72%) mostly presented with CD, which can affect any part of the gastrointestinal tract (Chi-Square test, p = 0.012). Consistently, while 64% of VEO-IBD patients exhibited colonic-only disease, older-onset patients showed an equal distribution between colonic-only and ileocolonic involvement. Notably, the panenteric phenotype, a hallmark of pediatric CD characterized by inflammation spanning the small bowel, large bowel, and upper gastrointestinal tract, with isolated ileal or colonic disease being less common,was more prevalent in VEO-IBD patients (20%) compared to those with older-onset IBD (9%) (Chi-Square test, p = 0.024).^43^ Although prior data indicate that the proportion of monogenic disorders with IBD-like presentations decreases with increasing age of onset, we observed a higher enrichment of monogenic IBD mutations among older-onset patients (50%) compared to those with VEO-IBD (10%) (Fisher’s exact test, p = 0.0015). Several factors may explain this unexpected enrichment. First, the rare variants identified in monogenic IBD genes among older-onset patients were not all confirmed to be necessarily disease-causing. Second, because this was a VEO-IBD–focused cohort, older-onset patients included were more likely to have been preselected based on features suggestive of monogenic disease. Third, emerging evidence indicates that monogenic IBD may be more common in older pediatric and adolescent patients than previously recognized.^44,45^ Since genetic testing is not routinely performed in older children and adults, clinicians should consider monogenic IBD more broadly, particularly in cases with suggestive features (e.g. extraintestinal comorbidities or poor responses to therapy), regardless of age at diagnosis. Further details on surgical interventions, monogenic mutations, extraintestinal symptoms, and medications for each donor are provided in **Supplementary Table 1.**

### IEOs Exhibit Reduced Long-Term Growth Capacity

Intestinal biopsies were collected from various anatomical locations in the small and large intestines of patients with VEO-IBD, older-onset pediatric IBD, and age-matched controls to generate organoids. Demographic profile of the collected intestinal biopsies from donors is presented in **Table 1**. In cases where multiple biopsies were taken from the same donor, detailed information for each biopsy is provided in **Supplementary Tables 1 and 2**.

Intestinal organoids were successfully generated from all groups within one week of expansion following a single passage from crypt culture. The initial organoid generation rates were comparable between IBD and control samples, regardless of disease onset time (VEO: 70%, Older onset: 91%, Control: 66%, Chi-Square test, p = 0.075) or IBD subtype (UC: 83%, CD: 63%, IBDU: 74%, Control: 66%, Chi-Square test, p = 0.1) **(Figures 1A-B)**. Additionally, no significant association was observed between organoid generation rates and the original intestinal location (duodenum: 63%, ileum: 78%, colon: 67%, rectum: 80%; Chi-Square test, p = 0.28) **(Figure 1C)** or inflammation status of the biopsy sample (Inflamed: 67%, Non-inflamed: 68%; Fisher’s exact test, p = 0.85) **(Figure 1D)**. However, a substantial loss of growth was observed across all groups during subsequent passaging and after freeze-thawing. Organoid lines were considered successfully expanded if they could be passaged at least three times post-thaw. While initial organoid generation was largely successful, the proportion of lines capable of sustained expansion was lower across all groups (VEO: 44%, Older onset: 39%, Control: 45%) **(Figure 1A)**, irrespective of IBD type, intestinal location, or inflammation status of the original biopsy **(Figures 1B-D)** (McNemar’s test, p < 0.05).

**Figure 1.**
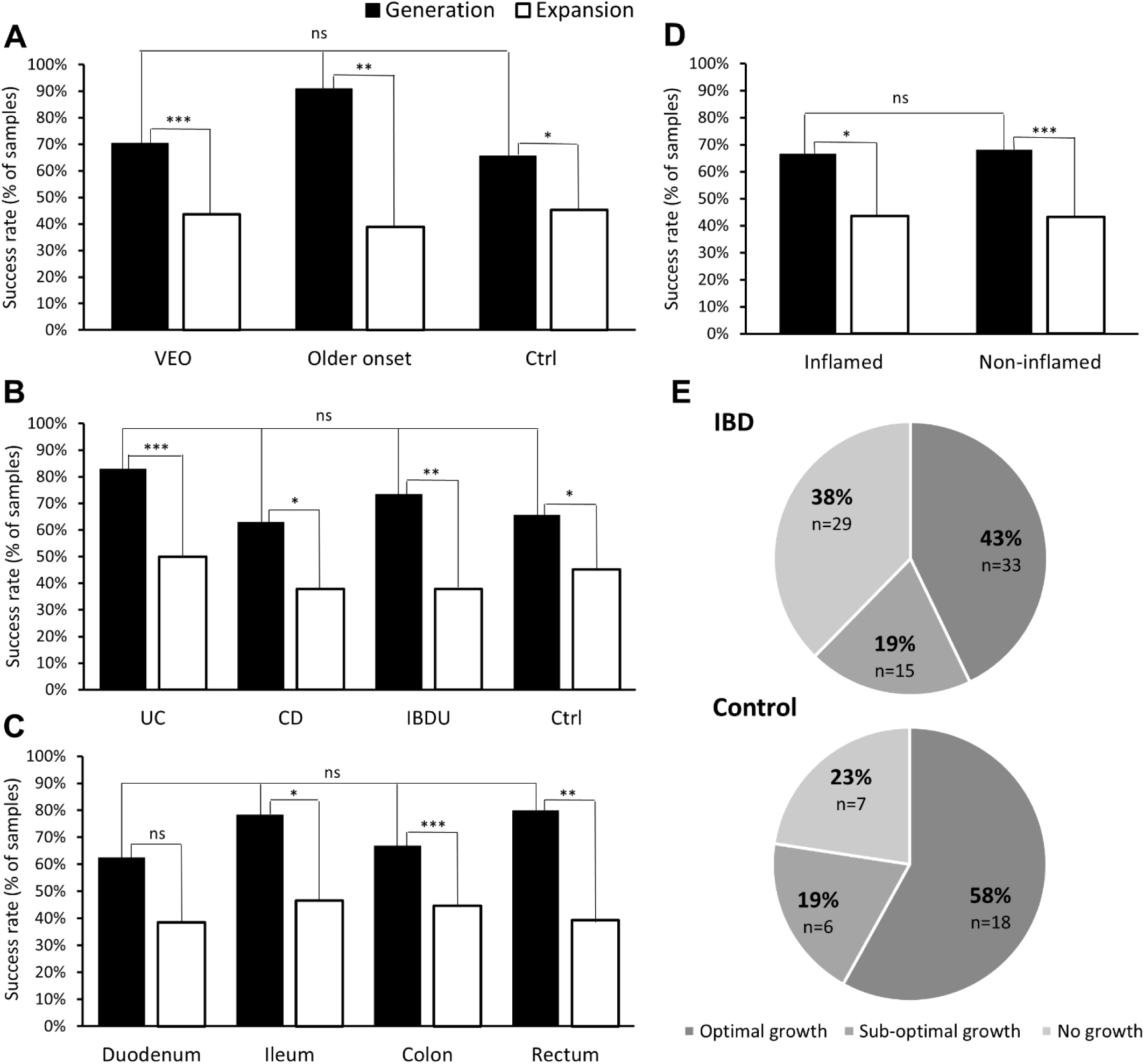
Intestinal organoids exhibit reduced long-term growth capacity. Bar graphs illustrate the success rates of organoid generation (black bars) and expansion (white bars) across different categories: **(A)** patient groups, sample sizes were: generation (VEO=111, older onset=22, control=64); expansion (VEO=94, older onset=18, control=53) **(B)** disease subtypes, sample sizes were: generation (UC=53, CD=46, IBDU=34, control=64); expansion (UC=46, CD=37, IBDU=29, control=53) **(C)** intestinal locations, accumulative sample sizes from both IBD and ctrl were: generation (Duodenum=16, Ileum=37, Colon=109, Rectum=35); expansion (Duodenum=13, Ileum=30, Colon=94, Rectum=28), and **(D)** inflammation status, accumulative sample sizes from both IBD and ctrl were: generation (Inflamed=48, Non-inflamed=133); expansion (Inflamed=39, Non-inflamed=97). Organoid lines where culture expansion was not assessed were excluded from the expansion analysis. Successful organoid generation is defined as the formation of growing organoid cultures within one week after a single passage from crypt culture, while successful organoid expansion is characterized by achieving more than three consecutive passages after freeze-thawing. Statistical comparisons of organoid generation rates between groups were conducted using the ChiSquare test, except for (D), which was analyzed using Fisher’s exact test. The significance of the decline from generation to expansion across different categories was assessed using McNemar’s test (*p <.05, **p <.01, ***p <.001). **(E)** Pie charts illustrate the distribution of organoid culture expansion outcomes in IBD (top) and control (bottom) samples, including only organoids for which expansion was assessed. expansion outcomes are categorized as follows: (1) Optimal growth (weekly expansion ratio >1:2), (2) Suboptimal growth (intermittent rather than consistent weekly expansion), and (3) No growth. Percentages and sample sizes are indicated for each group. No statistically significant difference was observed in the distribution of expansion outcomes between IBD and control samples (Chi-Square test, p = 0.27).

Further analysis of culture expansion revealed that 43% of established IBD organoid lines maintained optimal growth (consistently achieving a weekly expansion ratio >1:2) after freeze-thawing, compared to 58% of control lines. Suboptimal growth, characterized by intermittent rather than consistent weekly expansion, occurred at similar rates in both groups. However, a greater proportion of IBD lines (38%) failed to expand entirely, compared to 23% of control lines **(Figure 1E)**. These differences in expansion outcome distribution between the IBD and control groups were not statistically significant (Chi-Square test, p = 0.27).

### IEOs Display Donor-Specific Morphological Variations

Morphology and confluency were monitored 15 days after seeding 10,000 single cells from a subset of organoids (IBD=14, Control=3; **Supplementary Table 1**). Regardless of disease status, organoids from different donors, even from the same anatomical location, varied in size, shape, and confluency **(Figure 2A)**. For instance, donor 97’s transverse colon organoids were more confluent than donor 95’s, despite both being from inflamed transverse colon areas in VEO-UC patients. Similarly, descending colon organoids from IBD donor 90 were larger and more confluent than those from control donor 91. Sigmoid organoids ranged from normal cystic (IBD donor 85) to aberrant, with collapsed structures (IBD donor 89) or indiscernible aggregates (Control donor 102); IBD donor 85’s cultures were more confluent than IBD 89’s. IBD rectal organoids also varied: donor 51’s were cystic, while donor 12’s were primarily aggregated. Overall, 9 of the 14 IBD lines (64%) displayed typical cystic morphology, while in the remaining lines, a mixed morphology was observed, featuring both cystic organoids and non-cystic structures, such as collapsed or aggregated forms **(Figure 2B)**.

**Figure 2.**
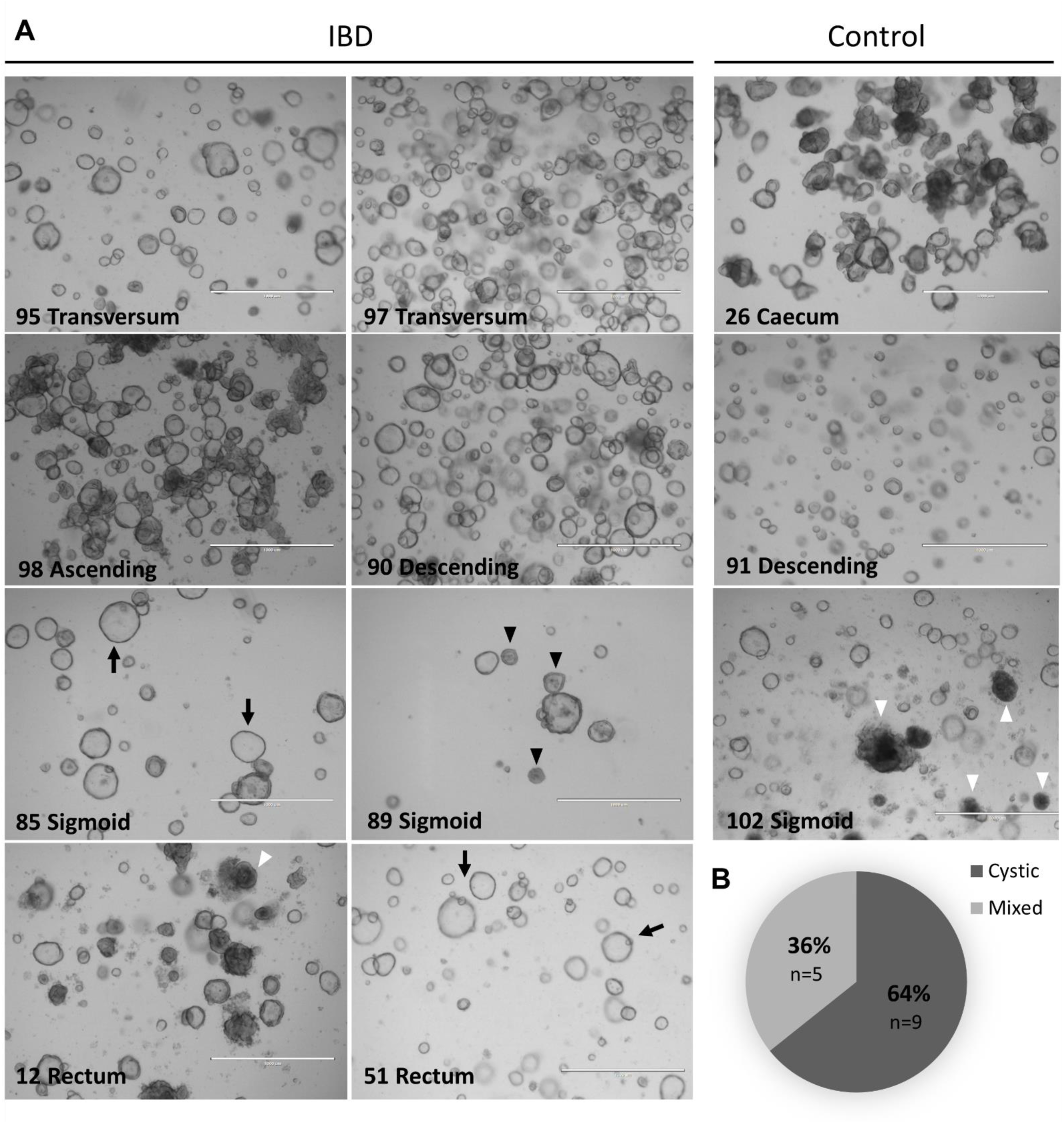
Morphological differences between organoids derived from IBD and Control Donors. **(A)** Representative bright field images of intestinal organoid cultures derived from various colon regions of IBD patients and control donors, taken 15 days after seeding 10,000 single cells per well in a 24-well plate, cultured in expansion medium. Notably, regardless of disease status, organoids from different donors, assigned by numbers, even when derived from the same anatomical location, exhibit significant variability in confluency, size, and shape. The observed morphologies include cystic structures (black arrow) with swollen, circular organoids, collapsed structures (black arrowhead), characterized by densely packed, non-cystic formations and aggregated structures (white arrowhead) where distinct shapes could not be identified. Images were obtained using a stereomicroscope (EVOS) at 4X magnification. **(B)** The majority of lines (64%, n=9/14) exhibited typical cystic morphology, while a subset (36%, n=5/14) displayed a mixed morphology, characterized by the presence of both cystic and non-cystic (collapsed or aggregated) structures.

### Bacterial Stimulation Effectively Re-induced Inflammation and Elicited IBD-Specific Responses in IEOs

To identify patient-specific epithelial phenotypes and therapeutic targets, we generated a comprehensive transcriptomic dataset of 204 intestinal organoid samples from successfully expanded lines. These samples, cultured for 3–4 weeks after thawing from frozen stocks, were derived from 24 unique IBD patients and 15 control donors **(Supplementary Tables 1 and 2)**. Among the 24 patients, eight were identified with specific monogenic variants in IBD-associated genes such as *IL10RA*, *TTC7A*, *LRBA*, *STXBP2*, *BTK,* and *TRNT1* **(Supplementary Table 1)**.^12,46^

IBD organoids gradually lose their transcriptional inflammatory signatures over time in the absence of inflammatory stimuli.^28,47^ This makes them an ideal platform for identifying intrinsic epithelial-driven immune responses, independent of the original biopsy’s inflammatory state, which may be central to disease pathogenesis. To uncover abnormal responses in IBD organoids, RNA-seq was conducted both at baseline and following immunological challenge for 6 and 72 hours to simulate a pathologically relevant environment. The immunological challenge was conducted using a specific assay we previously developed.^48^ This assay effectively stimulates intestinal stem cells in organoids with an *E. coli* lysate when applied basolaterally, providing a controlled and diverse array of bacterial antigens.

Exposure to bacterial lysate effectively re-induced inflammation in both control and IBD organoids, as demonstrated by the upregulation of pro-inflammatory markers, including *CXCL8*, *NFKB2*, *IL-23A*, *BIRC3*, *TNFAIP1*, and *DUSP16* **(Figure 3A)**. This response was accompanied by the activation of key inflammatory signaling pathways, such as IL-17A, NF-κB, TNF, and cytokine-related pathways **(Figure 3B)**. At both baseline and post-stimulation, IBD and control organoids remained indistinguishable, while PCA analysis identified bacterial stimulation as the main driver of variation **(Figure 3C)**.

**Figure 3.**
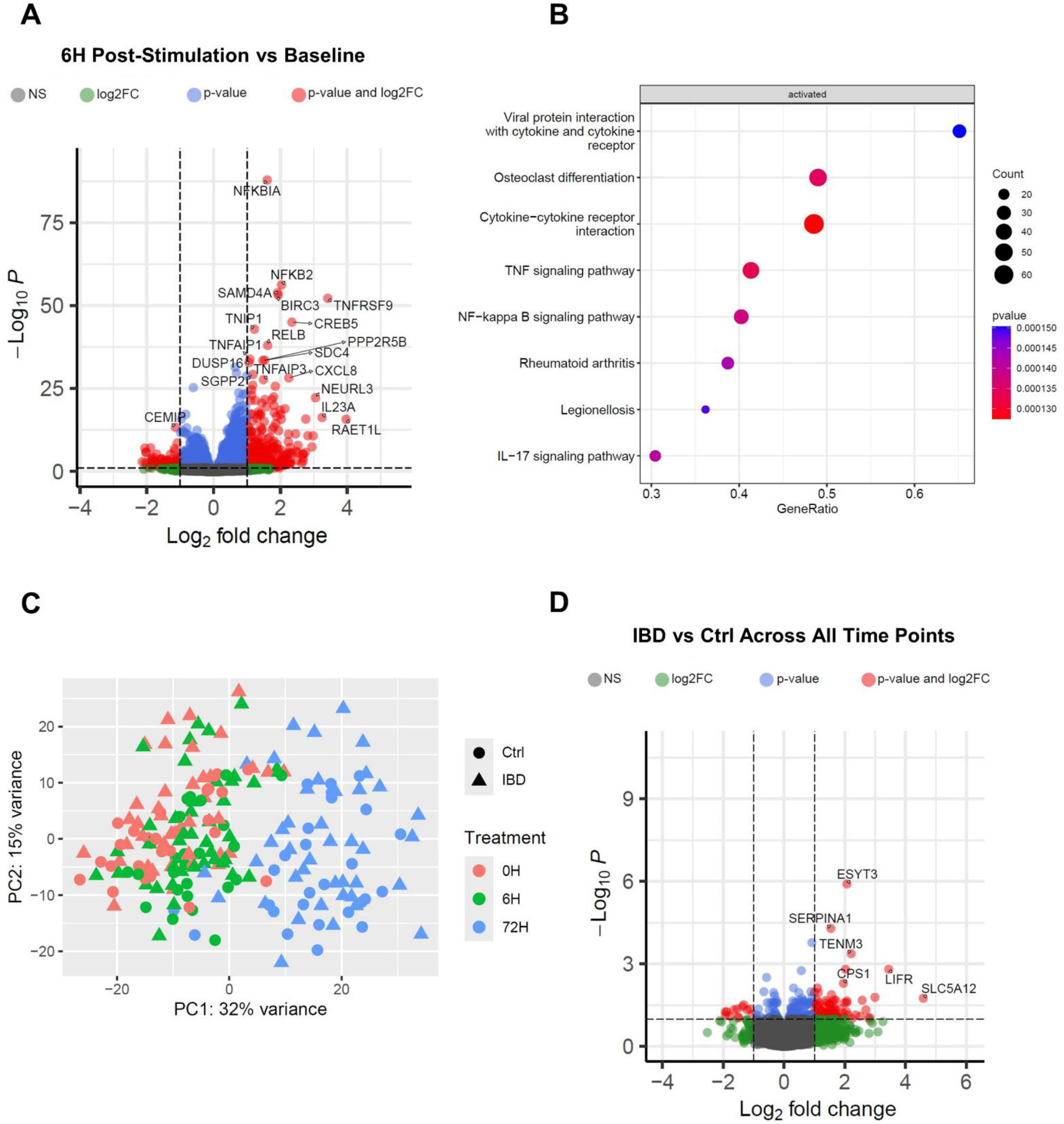
Bacterial stimulation re-induce inflammation in IEOs. **(A)** Differential gene expression analysis comparing 6H stimulated vs. baseline conditions in IBD and control organoids. The volcano plot highlights genes with significant up- or downregulation (adjusted p-value <0.01) (red), emphasizing the upregulation of key pro-inflammatory markers, including *CXCL8*, *NFKB2*, *IL-23A*, *BIRC3*, *TNFAIP1*, and *DUSP16*. **(B)** Pathway enrichment analysis of differentially expressed genes (DEGs) following 6H organoid stimulation in both IBD and control groups. The dot plot shows significantly activated inflammatory pathways, including IL-17, NF-κB, TNF, and cytokine-cytokine receptor interactions, with dot size representing gene count and color indicating statistical significance. **(C)** Principal Component Analysis (PCA) of organoid samples based on transcriptomic profiles. Stimulated (6H, 72H) and non-stimulated (0H) samples cluster separately, while IBD and control organoids remain largely indistinguishable at both baseline and post-stimulation. **(D)** Differential gene expression analysis comparing IBD vs. control organoids. The volcano plot highlights significantly DEGs (adjusted p-values <0.01) in IBD-derived organoids across all time points (0H, 6H, 72H), suggesting distinct transcriptional responses to bacterial stimulation.

However, gene expression profiles at all time points displayed differential changes in IBD-derived organoids as compared to controls, with notably upregulation of *SERPINA1*, *LIFR*, *ESYT3, CPS1*, *TENM3*, and *SLC5A12* **(Figure 3D)**. Among these, *SERPINA1* and *LIFR* are strongly linked to inflammation and immune responses, with implications in IBD.^49–52^ *SERPINA1*, which encodes α1-antitrypsin (AAT), protects tissues from inflammation caused by infection, injury, and inflammatory conditions like IBD.^49^ LIFR encodes receptor for the pleiotropic cytokine leukemia inhibitory factor (LIF). LIF is involved in various processes including immune responses in a context-dependent manner.^52^ While *ESYT3* shows emerging evidence of involvement in immune cell signaling, *CPS1* and *TENM3* are primarily linked to metabolic and neural functions, respectively, but may contribute to immune dysregulation.^53,54^ *SLC5A12*, encoding a transporter protein, is not directly related to immune processes.

### Network Analysis Reveals Subnetworks Associated with IEO Responses to Bacterial Stimulation and Various Genetic Mutations

The minimal overlap in affected genes across patients made it challenging to derive meaningful conclusions from individual gene expression profiles. To overcome this limitation, we applied a Weighted Gene Co-Expression Network Analysis (WGCNA) approach to the 192 transcriptional profiles, enabling us to better define and understand the complex biological processes underlying epithelial responses to bacterial stimulation and the impact of specific genotypes on these responses. This method, previously used to correlate gene modules with clinical parameters in pediatric IBD, enables the identification of gene clusters (modules) with correlated patterns, which can be summarized through module eigengenes and associated with phenotypes (e.g. genetic mutations) of interest to better understand their impact.^55^ A WGCNA was constructed across the entire cohort of control and IBD intestinal organoid samples (see Methods), including samples from different time points with the rationale of capturing dynamic transcriptional profiles across various states that better mimic IBD as it includes quiescent (0 hour, unstimulated) as well as active disease (6 hours of stimulation) and remission disease states (72 hours of stimulation). The signed organoid WGCNA identified 12 distinct co-expressed gene modules of varying sizes **(Table 2, Supplementary Table 4)**.

**Table 2.**
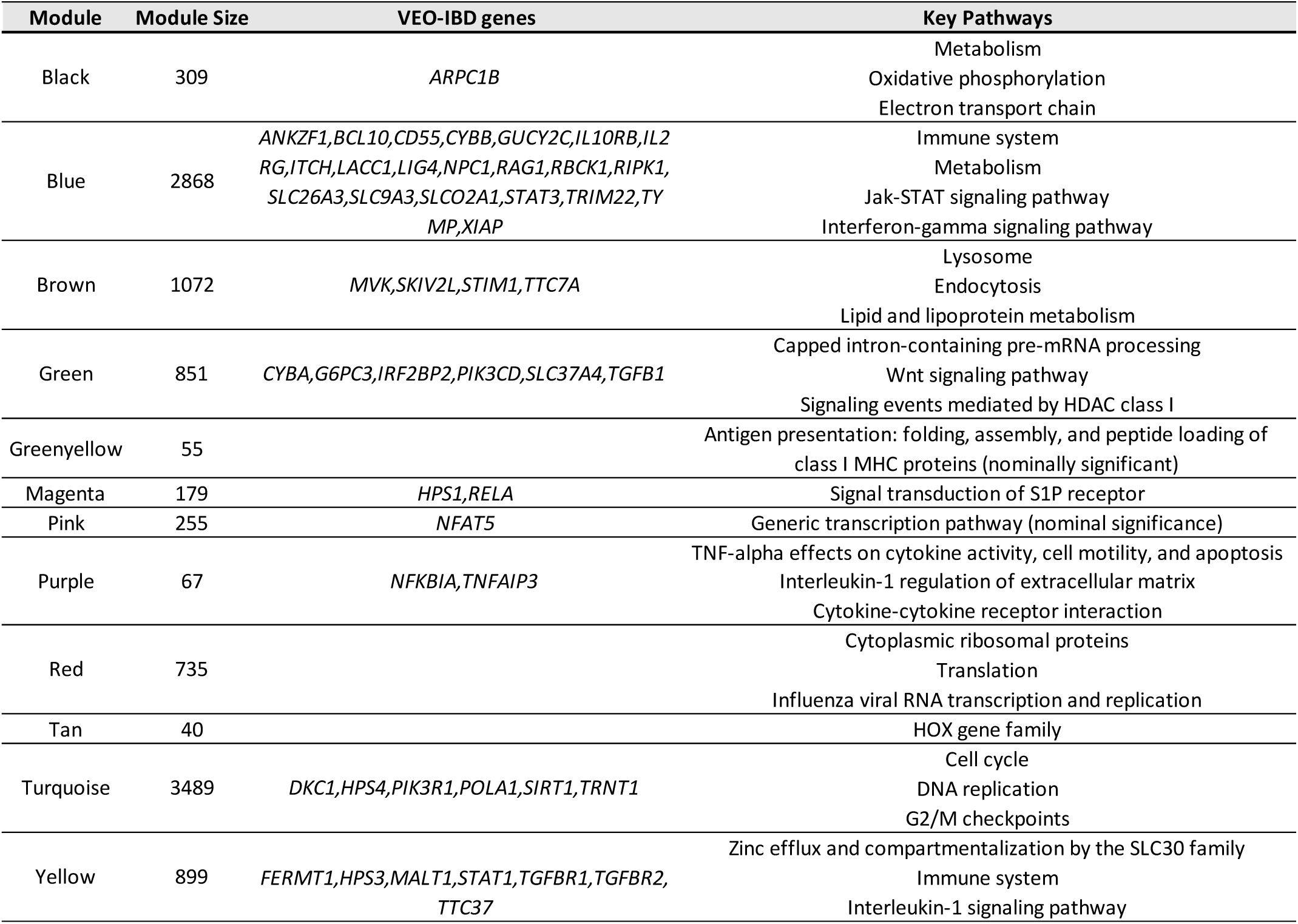
Key pathways and intersecting VEO-IBD genes across modules.

Modules of co-expressed genes often indicate shared functions. The results of pathway and gene enrichment analyses, using BioPlanet database terms and IBD-related genes respectively, are presented in **Supplementary Tables 5 and 6**. **Table 2** presents a selection of the top enriched biological pathways along with intersecting genes associated with VEO-IBD across various modules. Notably, all modules—except for greenyellow, red, and tan—contain genes known to be linked to VEO-IBD.

To identify potential connections between gene expression patterns and organoid-related phenotypes or genotypes, we analyzed the association of each gene module eigengene (a representative gene expression profile) with various traits, including bacterial exposure time, disease status (Ctrl vs IBD), and specific genotypes. Module-trait correlation analysis revealed varying degrees of association between gene module expression and these factors. After adjusting the transcriptional profiles of organoid samples for potential confounding effects due to differences in intestinal location (Location_SIvsLI) **(See QC plots in Supplementary Figure 1)**, minimal associations with this factor were observed. Instead, bacterial stimulation (Exposure time_inAll) was identified as a

a significant driver of gene module expression variation across the dataset **(Figure 4A)**. Bacterial stimulation significantly upregulated the purple module in both IBD and control organoids, peaking at 6 hours **(Figure 4A and Supplementary Figure 3A)**. Enriched for pro-inflammatory pathway terms related to TNF-α and IL-1 signaling **(Table 2)**, this module encompasses all six pro-inflammatory IBD genes (*CXCL8*, *NFKB2*, *IL-23A*, *BIRC3*, *TNFAIP1*, and *DUSP16*) previously identified by differential gene expression analysis (Figure 3A).^56–58^ Their co-expression within this module **(Supplementary Table 4)** strongly confirms the induction of an inflammatory response upon bacterial exposure. Furthermore, bacterial stimulation activated the blue module, notably enriched for IBD-driving inflammatory pathways like JAK-STAT and IFN-gamma signaling **(Table 2).**^59–61^ This module was also significantly enriched in VEO-IBD-associated genes, implicated in epithelial and immune defects, highlighting the profound influence of bacterial exposure on IBD-related transcriptional dynamics, extending beyond just inflammation **(Table 2 and Supplementary Table 5)**.

**Figure 4.**
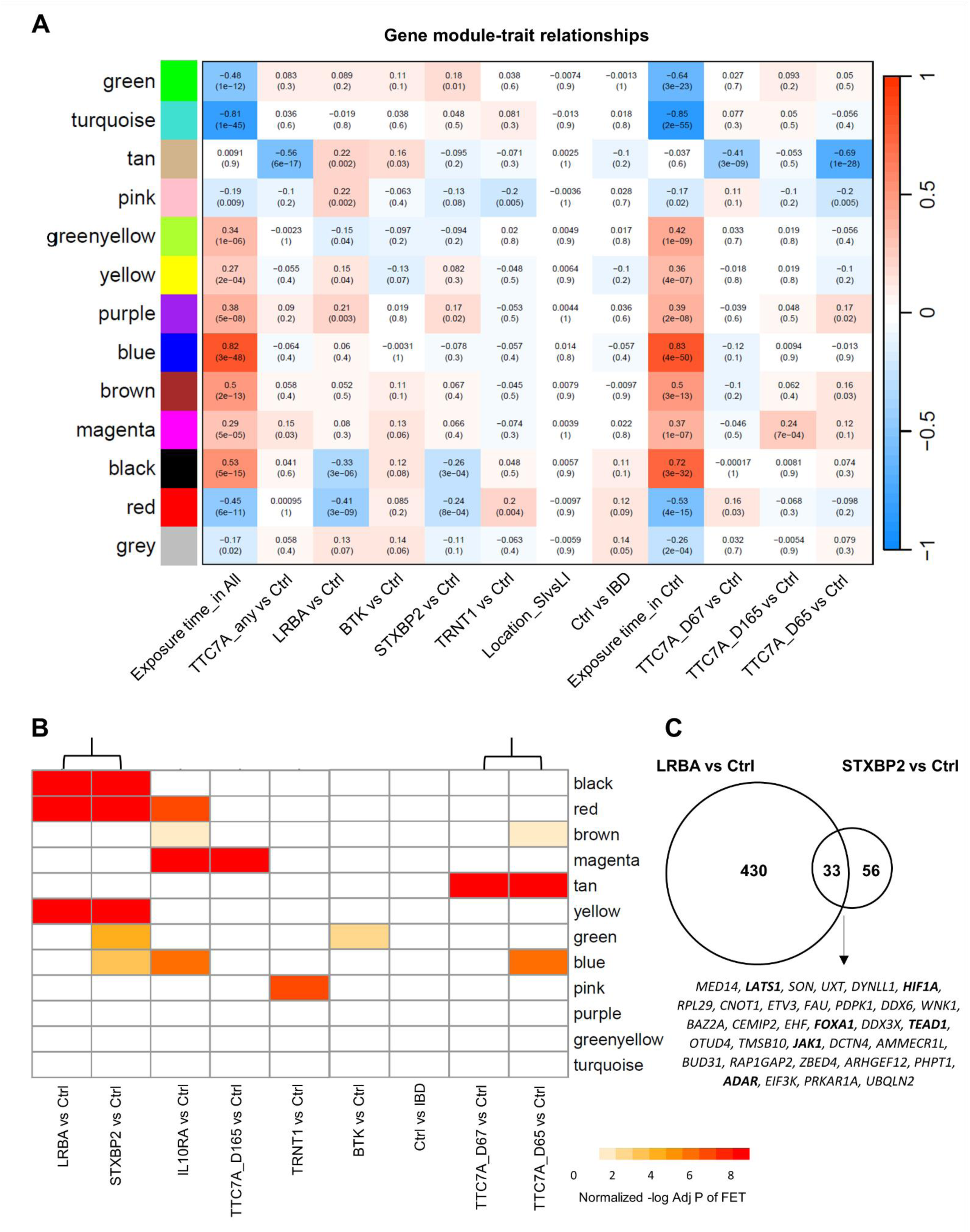
Network analysis of IEOs reveals transcriptional module associations with bacterial stimulation and specific genotypes. **(A)** The heatmap illustrates the correlations of the module eigengenes with traits based on signed Weighted Gene Co-Expression Network Analysis (WGCNA) performed on intestinal organoid transcriptomes from IBD and control samples (see Methods). Rows represent distinct modules of co-expressed genes, each labeled by a unique color on the left, as defined by WGCNA. Columns correspond to various traits and conditions, including disease status (Ctrl vs IBD), samples with genetic mutations vs control (e.g. distinct *TTC7A* variants from three different donors: D65, D67, and D165), Tissue location (small intestine vs large intestine, Location_SIvsLI), and exposure times to bacterial stimulation in control samples only (Exposure time_in Ctrl) and across all samples (Exposure time_in All). Each cell contains a correlation coefficient (-1 to 1) representing the relationship between a gene module and a given trait, with the p-value in parentheses indicating statistical significance. Red shades indicate positive association, with expression of the module eigengene being higher with the expression of the given trait. Blue shades indicate negative association, with lower expression of the module eigengene with the indicated trait. Bacterial stimulation exposure time is a continuous variable and has a strong association with expression of various modules. While general disease status (Ctrl vs IBD) does not show a strong correlation with most modules, several modules did associate with specific genotypes (e.g., *TTC7A*, *STXBP2*, *LRBA*). **(B)** Heatmap depicting the enrichment of genotype-based DEGs in WGCNA modules. Each column represents differentially expressed genes (DEGs) for a given genotype compared to the control (P-value < 0.05). The color intensity reflects the statistical significance of module enrichment (normalized-log10 adjusted P-value from Fisher’s Exact Test), with deeper red indicating stronger enrichment. **(C)** Venn diagram showing the overlap of DEGs (at AdjP-value < 0.05) between LRBA vs Ctrl and STXBP2 vs Ctrl. The numbers within each section indicate the count and proportion of unique and shared DEGs between the two comparisons.

We could not observe a significant association of overall disease status (Ctrl vs IBD) to module expression, possibly due to high inter-patient variability and the dominating effect of bacterial stimulation across the time points **(Figure 4A)**. However, we could observe differential module expression for some of the individual genotypes (notably *TTC7A*, *LRBA* and *STXBP2*) relative to controls. For example, we observed a striking effect of bacterial stimulation across the time points **(Figure 4A)**. However, we could negative association between the expression of the tan module in *TTC7A*-mutant samples, compared to control samples **(Figure 4A)**. The tan module consists of 40 genes, including *NOX1*, a gene that encodes NADPH oxidase 1 and has been linked to IBD.^5^ Pathway enrichment analysis, according to Bioplanet was uninformative, with no terms passing statistical threshold **(Supplementary Table 6)**, however, in reviewing the individual genes comprising the tan module, ∼19 genes of the 40 were annotated as homeobox (HOX) genes **(Supplementary Table 4)**. HOX genes are transcription factors that play a role in specifying the anterior-posterior axis during development and are important in demarcating intestinal subregions during gut development ^6^. Given the *TTC7A*-mutant samples were derived from different intestinal locations across three patients—D65 (colon), D67 (ileum), and D165 (duodenum)—each carrying distinct TTC7A variants **(Supplementary Table 1)**, we re-analyzed each TTC7A patient individually. When analyzing samples separately, the negative correlation of the tan module relative to controls appeared to be primarily driven by D65, partially by D67, and not at all by D165 and regardless of intestinal location or inflammatory stimulation (resting 0H and activated states 6H, 72H) **(Figure 4A and Supplementary Figure 3B)**. Enrichment analysis of differentially expressed genes (DEGs) from each of the three *TTC7A* genotypes (vs. controls), within WGCNA confirmed the *TTC7A* variant-specific association with the tan module **(Figure 4B, Supplementary Tables 7 and 8)**. In general, the D65 and D67 DEGs are enriched in similar modules as compared to D165 **(Figure 4B)**. These distinct patterns of module enrichment across organoids with different *TTC7A* mutations suggest that, despite affecting the same gene, different *TTC7A* variants drive heterogeneous transcriptional changes in epithelial organoids, depending on the specific mutation, in line with our previous observations in cell lines ^7^.

Interestingly, we observed similar module enrichment patterns for *STXBP2* and *LRBA* mutations when compared to controls. **(Figure 4B)**. DEGs associated with each genotype share significant enrichment in the black, red, and yellow modules **(Figure 4B and Supplementary Table 8)**. The expression of the black and red modules shows a negative association, whereas the yellow module exhibits a positive association in STXBP2 and LRBA samples compared to controls **(Figure 4A)**. When comparing genotype-associated DEGs pairwise, the strongest enrichment of LRBA-specific DEGs was observed with STXBP2-associated DEGs, showing the highest odd’s ratio (>6.0), despite significant enrichment also being detected for DEGs from several other genotypes **(Supplementary Table 9)**. The Venn diagram in **Figure 4C**, shows the overlap in molecular signatures between *LRBA* and *STXBP2* genotypes, according to DEGs at an adjusted p value <0.05. Among the intersecting genes are several key regulators of epithelial homeostasis, including: *LATS1* and *TEAD1*, which are core components of the Hippo signaling pathway, regulating intestinal epithelial regeneration and homeostasis ^8^; *HIF1A* and *JAK1*, respectively involved in hypoxic and inflammatory responses in the gut, both relevant to IBD ^9–11^; *ADAR*, which plays a role in RNA editing and is linked to immune regulation and gut epithelial homeostasis ^12,13^; and *FOXA1*, responsible for regulating intestinal epithelial cell (IEC) differentiation ^13^. Overall, this shared RNA expression pattern suggests that these mutations may contribute to disease pathogenesis through common pathways. These include increased expression of genes associated with ‘Zinc efflux and compartmentalization by the SLC30 family’, ‘Immune system’, and ‘Interleukin-1 signaling pathway’ all enriched in the yellow module, as well as reduced expression of genes in the black and red modules, which are respectively associated with ‘oxidative phosphorylation’ and ‘mRNA translation’ **(Table 2)**.

## Discussion

Despite the critical role of intestinal barrier dysfunction in IBD onset and persistence, a significant gap exists in targeted research and treatments addressing these epithelial components. To bridge this knowledge gap, we utilized patient-derived intestinal epithelial organoids (IEOs) from a pediatric IBD cohort and controls. Our study examined how inherent epithelial defects, including those driven by monogenic factors, contribute to pediatric IBD pathology and may serve as potential personalized therapeutic targets. Our analysis revealed transcriptional perturbations associated with bacterial antigen exposure and specific monogenic IBD factors.

Consistent with previous studies, we observed comparable organoid generation success rates between IBD (74%) and non-IBD control (66%) samples.^24,71^ However, while another study reported a lower IBD organoid generation rate compared to controls, their assessment differed methodologically, defining’normal growth’ as ≥50% crypt-to-organoid conversion within 24 hours post-plating, rather than monitoring growth after a single passage from crypt culture as in our study and the aforementioned consistent studies.^72^ Despite successful initial organoid generation, sustained growth decreased substantially after freeze-thawing and passaging in both IBD and control groups, independent of IBD type, intestinal location, or inflammation status of the original biopsy. This reduction can stem from a combination of factors including but not limited to: cell death during passaging due to disruption of cell-cell contacts and impaired Notch signaling, freeze-thawing-induced cellular damage, and potential effects of suboptimal expansion conditions (50% WCM, 10% NCM both derived from in-house cell lines) for maintaining intestinal stem cells undifferentiated.^48,73^

Consistent with previous studies showing that inflammation can be re-induced in both UC and non-IBD colon organoids using pro-inflammatory cytokines, we found that longterm cultured organoids effectively respond to E. coli lysate (containing diverse bacterial antigens, including LPS) by activating pro-inflammatory pathways and markers, regardless of disease status, origin, or mutation profile.^28^ Furthermore, while several genes (*SERPINA1, LIFR, ESYT3, CPS1, TENM3, and SLC5A12*) were differentially expressed between IBD and control samples after bacterial stimulation, only *SERPINA1* and *LIFR* have thus far been linked to IBD.

*SERPINA1*, a serpin gene encoding AAT, possesses anti-inflammatory and immunomodulatory properties relevant to pulmonary, autoimmune, and infectious diseases.^49^ Although primarily produced by hepatocytes, minor amount of SERPINA1 is also synthesized by immune, intestinal and bronchial epithelial cells.^49^ In active UC, *SERPINA1* is a hub gene for autophagy, with increased mucosal levels correlating with disease activity and reduced levels observed upon treatment.^50^ Multi-trait analysis of the IBD-nutrition-osteoporosis causal pathway identified *SERPINA1* as a deleterious variant in the European population.^74^ C-36 peptide, a degradation product of AAT, modulates human monocyte activation through LPS signaling pathways.^75^ The observed *SERPINA1* upregulation in our IBD organoids following bacterial lysate (including LPS) stimulation raises the possibility of a similar functional role in intestinal cells.

LIF, a pleiotropic cytokine of the IL-6 superfamily, has paradoxical pro-and anti-inflammatory effects depending on the different cells and diseases.^76^ However, studies suggest an anti-inflammatory role for LIF in UC, where it is upregulated in colon tissues, primary IECs, colitis mouse models, and LPS-treated Caco-2 cells.^51,77^ In a colitis mouse model, microbiota dysregulation induces LIF secretion by IECs restricting inflammation by inhibiting pathogenic T helper 17 differentiation and promoting epithelial proliferation via STAT4/STAT3 signaling.^78^ Suppression of LIF via microRNAs exacerbates epithelial inflammation in vitro and in vivo, while recombinant LIF restores microbiome homeostasis and mitigates induced colitis in mouse models, highlighting its therapeutic potential.^77,78^ In our IBD organoids, LIFR is upregulated without a corresponding increase in LIF, suggesting dysregulated LIF/LIFR signaling. This may reflect an attempt to enhance LIF sensitivity, compensate for downstream defects, or indicate ligand-independent activation.^79^ Given the context-dependent nature of LIF signaling and the heterogeneity of our IBD cohort (UC, CD, and IBDU), this pattern differs from previous studies that focused on UC alone and used different models than organoids. Our findings support targeting the LIF/LIFR pathway as a potential IBD therapy. However, further research is needed to fully understand its role and develop effective treatments.

In our co-expressed network analysis (WGCNA), we could not detect a significant association between IBD group as a whole and modules of co-expressed genes based on the entire dataset of samples. Transcriptional variation appeared primarily driven by bacterial stimulation, not disease status. This may be due, in part, to the limited sample size in our study, the heterogeneity of IBD patients and mechanisms underlying their disease as well as analyzing the cohort as a whole instead of within different time points or based on genes differential response to stimulation (e.g. change in expression at 6 hr vs 0 hr). However, this is an ongoing effort, and we aim to include more patients and different analytical approaches in future analyses to address these limitations. Previous studies comparing the transcriptomes of IBD and non-IBD organoids have reported both consistent and inconsistent findings with our study.^22,24,28,55,80,81^ These discrepancies may stem from various factors known to influence transcriptional variation in intestinal organoids, including media composition, sample size, intestinal location, differentiation status, donor variability, technical variations in culture conditions (batch-to-batch variations in conditioned media, incubator conditions, passage rate, and media change intervals), and differences in sequencing platforms.^22,82^

In contrast to overall disease status, specific gene mutations showed stronger associations with module expression. Notably, two of the three distinct *TTC7A* variants were uniquely linked to reduced expression of a gene module in which nearly half of the genes are annotated as HOX transcription factors. These HOX genes help guide gut axis patterning and shape the structure of the developing intestine.^63^ Furthermore, colon organoids from two patients with *LRBA* and *STXBP2* mutations shared a molecular phenotype characterized by activation of IL-1 signaling and pathways involved in intracellular zinc distribution via SLC30 transporters. This was accompanied by a negative association with modules enriched for oxidative phosphorylation and protein translation. Excessive IL-1 production may damage IECs.^83^ Additionally, disrupted zinc homeostasis, commonly linked to dysfunctional SLC30 transporters, directly impairs intestinal barrier integrity by affecting tight and adherens junctions.^84^ This improper zinc handling also compromises antioxidant defenses, fostering oxidative stress, a common feature in IBD, and disrupts normal epithelial proliferation, differentiation, and repair.^85,86^ Such shared epithelial phenotypes highlight the potential for patient stratification based on their molecular profiles.

Mutations in *STXBP2*, lead to familial Hemophagocytic Lymphohistiocytosis type 5 (FHL5), a rare hyper-inflammatory immune deficiency with many patients also presenting with gastrointestinal disorders such as IBD, severe diarrhea, and microvillus inclusion disorder.^87–94^ Importantly, IBD symptoms often persist even after successful hematopoietic stem cell transplantation (HSCT), highlighting STXBP2’s non-immune role in IBD pathogenesis.^88^ STXBP2 is essential for SNARE-mediated membrane fusion, apical cargo trafficking, and exocytosis in IECs, regulating the recycling and localization of apical membrane proteins crucial for intestinal polarity.^92,95,96^ While *LRBA* is a commonly reported monogenic IBD gene, its precise function in IBD pathogenesis remains

unclear.^44,97–103^ Anti-TNFα therapies are ineffective in *LRBA* deficiency, though HSCT can be beneficial.^44,46^ LRBA, a large intracellular protein, likely functions as an adaptor in vesicle regulation and lysosomal processes, including autophagy.^97,104,105^ LRBA is known to play a role in intracellular trafficking within immune cells.^98^ Given STXBP2’s essential role in intestinal epithelial cell trafficking, the similar transcriptional signatures_ including changes in zinc trafficking _ observed in both *LRBA* and *STXBP2* organoids could extend LRBA’s known trafficking function from immune cells to the epithelium. The shared transcriptional signatures identified in this study provide new insights into the pathogenesis of these two genotypes, that may be important for the development of future personalized, pathway-targeted therapies.

However, a potential limitation is that these intrinsic responses in IBD samples could partly reflect an epigenetic memory or’scarring’ from earlier inflammatory challenges, rather than purely a pre-existing epithelial dysfunction. This epigenetic imprinting might influence IBD-derived organoids’ long-term transcriptional behavior and their response to new stimuli. Our findings also raise key questions regarding the molecular mechanisms and functional consequences of the genotype-specific expression changes in the epithelium. Future studies using genome editing to correct or introduce these mutations in intestinal organoids could provide definitive conclusions about the specific functions of these genes in the epithelium and their contribution to IBD pathology.

## Conclusion

In summary, our study, using a large cohort of patient and control organoid samples, highlights the value of IEOs as a model for exploring epithelial phenotypes in pediatric IBD. We identified disease-specific epithelial disturbances and genotype-associated biological pathways, offering insights into epithelial dysfunction and monogenic causes while revealing potential targets for personalized therapies. Although our focus is on pediatric IBD, the shared phenotypic and pathogenic mechanisms between pediatric and adult-onset IBD suggest broader relevance for our findings.

## Acknowledgements

We thank all of the Boston Children’s Hospital, Klinikum der Universitat Munchen, and University Medical Center Utrecht patients and their families who consented and participated in this study as part of the Helmsley VEOIBD (www.VEOIBD.org) consortia and STEM study, as well as the health care professionals who care for these patients. We thank Utrecht Sequencing Facility (USEQ) for performing RNA sequencing.

## Author contributions

EN and MM contributed to study and experiment designs. HC and JHvE led the teams responsible for establishing the organoids. ZS and MHdV performed the experiments. EK and CA performed RNA-seq analysis. ZS performed organoid culture analysis. CA, EK, and ZS performed data interpretation. ZS wrote the manuscript. SBS, CK, LVC, DK, and MS provided patient biopsies and clinical data. EK, CA, MM, JHvE, LVC, and EN edited the manuscript and provided intellectual content. All of the authors fulfill the criteria for authorship and read and approved the final manuscript.

## Data availability statement

The patient dataset and processed RNA sequencing data will be made available upon publication.

