## Supplement for "A collection of patient-derived intestinal organoid lines reveals epithelial phenotypes associated with genetic drivers of pediatric inflammatory bowel disease"

### Supplementary Materials

#### Supplementary Table 1. Characteristics of IBD Donors and Derived Organoid Lines

| Donor ID | Dx | Age at Dx | Age at SC | Gender | Disease location | Monogenic mutation[a] | P. Inv | Stricturing | Surgical history | Medication | Extraintestinal comorbidities | Tissue | Inflamm. status | Org. Gen | Org. Exp | RNA seq[b] | Morphology[c] |
|---|---|---|---|---|---|---|---|---|---|---|---|---|---|---|---|---|---|
| D001 | IBDU | 1 | 5 | M | Unk | None | Yes | Unk | Colectomy | Unk | Unk | Ile | Unk | Yes | Opt | Yes | |
| D002 | IBDU | 1 | 10 | F | Pancolitis | None | Yes | Yes | Ileostomy, colectomy | Unk | Unk | Rec | Infl | Yes | Unk | No | |
| D003 | IBDU | 2 | 4 | M | Unk | None | Unk | Unk | Unk | Unk | Unk | Ile | Unk | Yes | No | No | Cystic |
| | | | | | | | | | | | | Col | Unk | Yes | Opt | Yes | |
| | | | | | | | | | | | | Rec | Unk | Yes | Sub-opt | Yes | |
| D004 | IBDU | 15 | 15 | M | Pancolitis | None | No | Unk | None | Omeprazole | None | AC | Non-infl | Yes | No | No | |
| | | | | | | | | | | | | TC | Non-infl | Yes | No | No | |
| | | | | | | | | | | | | DC | Non-infl | Yes | No | No | |
| D006 | UC | 3.3 | 6 | F | Pancolitis | None | No | N/A | None | Balsalazide, azathioprine, omeprazole, | None | Ile | Non-infl | Yes | Unk | No | |
| | | | | | | | | | | | | Cae | Non-infl | Yes | Unk | No | |
| | | | | | | | | | | | | Rec | Non-infl | Yes | Sub-opt | No | |
| D007 | UC | 3.5 | 5 | F | Colonic | STXBP2 | No | N/A | Colectomy-ileostomy, pouch surgery, ileostomy closure and J-pouch | Prednisone | Fever without source | Ile | Non-infl | Yes | No | No | |
| | | | | | | | | | | | | Cae | Non-infl | Yes | Opt | Yes | |
| | | | | | | | | | | | | Rec | Non-infl | Yes | Sub-opt | Yes | |
| D009 | UC | 4.8 | 5 | M | Pancolitis | None | No | N/A | None | Amoxicillin, metronidazole, acetaminophen, ranitidine | None | Ile | Non-infl | No | N/A | No | |
| | | | | | | | | | | | | Cae | Non-infl | Yes | Opt | Yes (N=2) | |
| | | | | | | | | | | | | Rec | Non-infl | Yes | No | No | Non-cystic |
| D011 | CD | 0.4 | 2 | F | Colonic | IL10RA | Yes | No | Colostomy | None | None | Duo | Non-infl | Yes | Opt | Yes | |
| | | | | | | | | | | | | Ile | Non-infl | Yes | Opt | Yes | |
| | | | | | | | | | | | | Rec | Non-infl | Yes | Unk | No | |
| D012 | UC | 2 | 2 | F | Pancolitis | None | No | N/A | None | None | None | Duo | Non-infl | Yes | Opt | Yes | |
| | | | | | | | | | | | | Ile | Non-infl | Yes | Unk | No | |
| | | | | | | | | | | | | Cae | Non-infl | Yes | Opt | Yes | |
| | | | | | | | | | | | | Rec | Non-infl | Yes | Opt | Yes | Non-cystic |
| D014 | UC | 3 | 3 | M | Pancolitis | None | No | N/A | Colectomy-ileostomy, pouch surgery | None | None | Duo | Infl | No | N/A | No | |
| | | | | | | | | | | | | Ile | Infl | Yes | Opt | Yes (N=2) | |
| | | | | | | | | | | | | Sig | Infl | Yes | Opt | Yes | |
| D015 | CD | 6.5 | 16 | M | Ileocolonic | BTK | No | No | None | Intravenous immunoglobulin | X-linked Agammaglobulinemia | Duo | Unk | Yes | No | No | |
| | | | | | | | | | | | | Cae | Unk | No | N/A | No | |
| | | | | | | | | | | | | AC | Unk | Yes | Sub-opt | Yes | |
| D016 | UC | 1.2 | 2 | F | Pancolitis | None | No | N/A | None | Sulfasalazine | None | Duo | Non-infl | Yes | Opt | Yes | |
| | | | | | | | | | | | | Ile | Non-infl | Yes | Opt | Yes | |
| | | | | | | | | | | | | AC | Non-infl | Yes | Opt | Yes | |

#### Supplementary Table 1. Continued

| | | | | | | | | | | | | | | | | | |
|---|---|---|---|---|---|---|---|---|---|---|---|---|---|---|---|---|---|
| | | | | | | | | | | | | Rec | Infl | Yes | Unk | No | |
| D017 | UC | 0.2 | 2 | F | Colonic | None | No | N/A | None | Sulfasalazine, folic Acid | None | Cae | Unk | Yes | Unk | No | |
| | | | | | | | | | | | | Col | Unk | Yes | Opt | Yes (N=2) | |
| | | | | | | | | | | | | Rec | Unk | Yes | No | No | |
| D018 | CD | 18 | 19 | F | Colonic | LRBA | No | No | None | Unk | None | Cae | Infl | Yes | Opt | Yes | |
| | | | | | | | | | | | | Col | Infl | Yes | No | No | |
| D021 | IBDU | 0.9 | 1 | F | Colonic | None | No | No | None | Sulfasalazine, ferrous sulfate, folic acid, polyethylene glycol | None | Duo | Non-infl | No | N/A | No | |
| | | | | | | | | | | | | Ile | Non-infl | No | N/A | No | |
| | | | | | | | | | | | | Cae | Non-infl | No | N/A | No | |
| | | | | | | | | | | | | Rec | Non-infl | Yes | Unk | No | |
| D023 | IBDU | 0.8 | 1 | M | Pancolitis | None | No | Unk | None | Mesalamine, prednisone, pantoprazole | None | AC | Non-infl | Yes | No | No | |
| D029 | IBDU | 4.6 | 8 | F | Colonic | None | No | No | None | Sulfasalazine, prednisone | None | Ile | Non-infl | Yes | Sub-opt | Yes | |
| | | | | | | | | | | | | TC | Infl | Yes | Unk | No | |
| D037 | CD | 2.1 | 13 | F | Colonic | None | No | No | None | None | None | Col | Non-infl | Yes | Sub-opt | Yes | |
| D039 | CD | 7.1 | 6 | M | Ileocolonic | BTK | Yes | No | None | Mesalamine | X-linked Agammaglobulinemia | Ile | Unk | Yes | Opt | Yes (N=2) | |
| | | | | | | | | | | | | AC | Unk | Yes | Unk | No | |
| D040 | UC | 5.7 | 8 | F | Pancolitis | None | No | N/A | None | Claritin | None | AC | Infl | Yes | No | No | |
| D041 | UC | 4.7 | 8 | M | Colonic | None | No | N/A | None | None | None | Rec | Non-infl | Yes | Sub-opt | No | |
| D045 | CD | 7 | 7 | M | Ileocolonic | TRNT1 | No | No | None | None | None | Rec | Non-infl | Yes | Opt | Yes | |
| D046 | UC | 6.1 | 9 | F | Pancolitis | None | No | N/A | None | Mesalamine, vancomycin, budesonide, 6-MP | None | Rec | Infl | Yes | Sub-opt | No | |
| D047 | CD | 2 | 15 | F | Ileocolonic | None | No | Yes | Ileum resection, stricturoplasty | Infliximab, multivitamin, D3, ferrous sulfate, Miralax, eszopiclone | None | Ile | Infl | Yes | No | No | |
| | | | | | | | | | | | | TC | Non-infl | Yes | No | No | |
| D048 | UC | 5 | 11 | F | Colonic | None | No | N/A | None | Mesalamine, D3, multivitamin | None | Cae | Non-infl | Yes | Opt | Yes | |
| D049 | UC | 0.2 | 5 | F | Colonic | None | No | N/A | None | Sulfasalazine, folic Acid | None | Ile | Unk | Yes | No | No | |
| D050 | CD | 6.3 | 6.3 | F | Colonic | None | No | No | None | None | None | Cae | Infl | Yes | Opt | Yes | |
| D051 | CD | 5.3 | 8 | F | Ileocolonic | None | Yes | No | None | Ciprofloxacin, metronidazole, methylprednisone, infliximab, methotrexate | None | Rec | Infl | Yes | Opt | Yes | Cystic |
| D052 | UC | 2.4 | 2.4 | M | Colonic | None | No | N/A | None | Fer-In-Sol, multivitamins | None | Sig | Infl | Yes | Opt | Yes | |
| D053 | UC | 5 | 14 | M | Pancolitis | Unk | No | N/A | None | Mesalamine, budesonide, papaya | None | Ile | Non-infl | Yes | Sub-opt | No | |
| | | | | | | | | | | | | Rec | Non-infl | Yes | No | No | |

#### Supplementary Table 1. Continued

| | | | | | | | | | | | | | | | | | |
|---|---|---|---|---|---|---|---|---|---|---|---|---|---|---|---|---|---|
| | | | | | | | | | | enzyme extract, Ventolin | | | | | | | |
| D054 | UC | 5.7 | 6 | F | Pancolitis | None | No | N/A | None | Vancomycin, methylprednisone, folic acid, omeprazole, multivitamin | None | Rec | Infl | Yes | Sub-opt | No | |
| D055 | UC | 4.7 | 8 | F | Pancolitis | None | No | N/A | None | Infliximab , balsalazide | None | Ile | Non-infl | Yes | No | No | |
| | | | | | | | | | | | | Col | Non-infl | Yes | Unk | No | |
| D059 | CD | 0.5 | 9 | M | Colonic | None | Yes | No | None | Smoflipid Intravenous Emulsion, alteplase, GoLytely, ascorbic acid, emollients, ergocalciferol, ethambutol, famotidine, fluconazole, isoniazid, ketoconazole topical, loratadine, metronidazole, nitazoxanide, nystatin, pyrazinamide, pyridoxine, acetaminophen, Domeboro, ondansetron, | Fever | AC | Infl | No | N/A | No | |
| D060 | CD | 3.4 | 16 | M | Colonic | None | Yes | No | None | 6-MP, multivitamin, Tums Kids 750 | None | Ile | Non-infl | No | N/A | No | |
| | | | | | | | | | | | | Col | Non-infl | No | N/A | No | |
| D062 | CD | 7 | 7 | M | Ileocolonic | BTK | Yes | No | None | Mesalamine | X-linked Agammaglobulinemia | Ile | Non-infl | Yes | No | No | |
| | | | | | | | | | | | | Rec | Non-infl | Yes | No | No | |
| D063 | IBDU | 11 | 14 | F | Unk | None | No | Unk | Unk | Unk | Unk | Col | Unk | Yes | No | No | |
| | | | | | | | | | | | | Rec | Unk | Yes | No | No | |
| D065 | IBDU | 1 | 14 | M | Panenteric | TTC7A (Biallelic c.1027G>A; p.E343K and c.2405T>C; p.L802P) | No | Yes | Multiple balloon dilatations of the pylorus, jejunostomy, cholecystectomy | Unk | Acute Respiratory Distress Syndrome, allergic rhinoconjunctivitis, food allergies, growth hormone disorder | Col | Unk | Yes | Sub-opt | Yes (N=2) | |
| D066 | UC | 0.5 | 3 | M | Pancolitis | None | No | N/A | None | Mesalamine, D3 | None | Ile | Non-infl | Yes | No | No | |
| D067 | CD | 9 | 8 | M | Panenteric | TTC7A (Monoallelic c.1817A>G; p.K606R and c.2014T>C; p.S672P) | No | Yes | Exploratory laparotomy, ileostomy | Mesalamine, calcium carbonate, ergocalciferol, ethanol, heparin flush, pantoprazole, rifaximin, ursodiol, acetaminophen, aluminum sulfate-calcium acetate topical, nystatin topical, ondansetron | Food allergies, seborrheic dermatitis, steroid-induced diabetes mellitus, delayed puberty, short stature | Ile | Non-infl | Yes | Opt | Yes (N=2) | Non-cystic and fully collapsed |

#### Supplementary Table 1. Continued

| | | | | | | | | | | | | | | | | | |
|---|---|---|---|---|---|---|---|---|---|---|---|---|---|---|---|---|---|
| D068 | CD | 4.2 | 16 | F | Panenteric | Unk | Yes | Yes | Jejunum resection | Methotrexate, infliximab, folic acid, iron | None | Duo | Non-infl | No | N/A | No | |
| D069 | IBDU | 1.7 | 3 | M | Unk | None | Unk | Unk | Unk | Infliximab, tacrolimus, metronidazole, methylprednisone, proton pump inhibitor | Unk | Duo | Unk | Yes | Unk | No | |
| | | | | | | | | | | | | Ile | Unk | Yes | No | No | |
| D070 | CD | 1 | 15 | M | Unk | TTC37 | Unk | Unk | Unk | Unk | Tricho-Hepato-Enteric Syndrome | Rec | Unk | No | N/A | No | |
| D072 | UC | 1.4 | 2 | F | Colonic | None | No | N/A | None | None | Unk | Col | Non-infl | Yes | Opt | Yes | |
| D073 | UC | 5 | 9 | M | Pancolitis | None | No | N/A | None | Amoxicillin, metronidazole, acetaminophen, ranitidine | None | AC | Non-infl | No | N/A | No | |
| D074 | CD | 4.8 | 7 | M | Panenteric | None | Yes | No | None | Infliximab | None | Sig | Infl | Yes | Opt | Yes | |
| D075 | IBDU | 12 | 14 | M | Unk | None | No | Unk | Unk | Unk | Unk | TC | Unk | Yes | Unk | No | |
| D076 | CD | 11 | 11 | F | Unk | None | Unk | No | None | Azathioprine, atovaquone, fluticasone nasal, folic acid, sodium chloride nasal | None | Ile | Non-infl | Yes | No | No | |
| D078 | IBDU | 2 | Unk | M | Unk | None | No | Unk | Unk | Unk | Unk | Ile | Unk | Yes | No | No | |
| D083 | CD | 4.6 | 8 | F | Pancolitis | None | No | No | Ileal perforation at 5 months, status-post ileostomy with subsequent reanatsomosis | Sulfasalazine, ergocalciferol | None | Ile | Non-infl | Yes | Unk | No | |
| | | | | | | | | | | | | Col | Infl | No | N/A | No | |
| D085 | IBDU | 4.2 | 5 | F | Pancolitis | None | No | No | None | Sulfasalazine, vancomycin, ursodiol | Primary sclerosing cholangitis | Sig | Non-infl | Yes | Opt | Yes | Cystic |
| D087 | UC | 4.1 | 13 | M | Pancolitis | None | No | N/A | None | Sulfasalzine, 6-MP, hyoscyamine, folic acid | Unk | AC | Infl | No | N/A | No | |
| D089 | UC | 2 | 2.4 | F | Pancolitis | None | No | N/A | None | EpiPen as needed | None | Sig | Infl | Yes | Opt | Yes | Thick-walled |
| D090 | IBDU | 4 | 4 | M | UnK | None | No | UnK | UnK | UnK | Unk | DC | Infl | Yes | Opt | Yes | Cystic |
| D092 | IBDU | 1 | 8 | F | UnK | None | No | UnK | UnK | UnK | UnK | DC | Infl | Yes | Sub-opt | No | |
| D093 | CD | 4 | 13 | M | Pancolitis | None | Yes | Yes | None | Prednisone, vedolizumab | None | DC | Non-infl | Yes | No | No | |
| D095 | UC | 5 | 10 | M | Pancolitis | None | No | N/A | None | Acetaminophen, famotidine, fluoride topical, hydrocortisone topical, vedolizumab, ketoconazole topical, lorazepam, ondansetron, prednisolone, prednisone, triamcinolone topical | Arthopathy, oral apthous ulcers | TC | Infl | Yes | Opt | Yes | Cystic |

#### Supplementary Table 1. Continued

| | | | | | | | | | | | | | | | | | |
|---|---|---|---|---|---|---|---|---|---|---|---|---|---|---|---|---|---|
| D097 | UC | 1 | 1 | F | Pancolitis | None | No | N/A | Laparaoscopic subtotal colectomy and end ileostomy | Azathioprine, infliximab, ferrous sulfate, hydrocortisone, omeprazole, prednisolone | None | TC | Infl | Yes | Opt | Yes | Cystic |
| D098 | IBDU | 1 | 3 | M | Pancolitis | None | No | No | None | Ergocalciferol, folid acid, lactobacillus rhamnosus GG, multivitamin, sulfasalazine | None | AC | Infl | Yes | Opt | Yes | Non-cystic |
| D100 | CD | 0.8 | 1 | M | Panenteric | None | Yes | No | None | Canakinumab, multivitamin, vancomycin | Possible sinusitis | DC | Infl | No | N/A | No | |
| D104 | CD | 1 | 1 | M | Panenteric | None | No | No | None | Acetaminophen, albuterol, cholecalciferol, hyoscyamine, immune globulin subcutaneous, infliximab, meloxicam, multivitamin | None | Sig | Non-infl | No | N/A | No | |
| D106 | CD | 2 | 13 | M | Panenteric | None | No | No | Ileostomy | Motilium, Litican, Budenofalk, Zantac, morphine, Perfusalgan, Cefurim, Flagyl, SoluMedrol | Atopic dermatitis, secondary glaucoma, osteopenia, dyslipidemia | DC | Infl | No | N/A | No | |
| D108 | IBDU | Unk | Unk | M | Unk | None | Unk | Unk | Unk | Unk | Unk | Duo | Non-infl | No | N/A | No | |
| | | | | | | | | | | | | DC | Non-infl | Yes | Sub-opt | No | |
| D109 | CD | 4.8 | 5 | M | Colonic | Unk | Yes | No | None | None | None | Rec | Infl | Yes | No | No | |
| D112 | UC | 5.7 | 13 | M | Colonic | None | No | N/A | None | Mesalamine | None | TC | Non-infl | Yes | No | No | |
| D113 | UC | 2 | 8 | M | Colonic | None | No | N/A | None | Sulfasalazine, azathioprine | None | DC | Infl | Yes | No | No | |
| D115 | CD | 4 | 17 | F | Ileocolonic | None | Yes | No | None | Stelara, sulfasalazine, methotrexate | Inflammatory arthritis, vulvar cutaneous Crohn's, erythema nodosum | DC | Non-infl | Yes | Sub-opt | No | |
| D116 | CD | 2.9 | 18 | M | Panenteric | None | Yes | No | None | 6-MP | None | TC | Non-infl | No | N/A | No | |
| D117 | CD | 5 | 17 | M | Ileocolonic | BTK | No | Yes | Ileocecectomy and small bowel resection, diverting ileostomy creation and lysis of adhesions, right hemicolectomy, and ostomy takedown, prolapse repairs, | Infliximab, Hizentra, topical creams (incl. tacrolimus) | Psoriasis, X-linked Agammaglobulinemia | DC | Non-infl | Yes | Sub-opt | No | Cystic |

#### Supplementary Table 1. Continued

| | | | | | | | | | | | | | | | | | |
|---|---|---|---|---|---|---|---|---|---|---|---|---|---|---|---|---|---|
| | | | | | | | | | small bowel resection | | | | | | | | |
| D119 | UC | 4 | 11 | F | Pancolitis | None | No | N/A | Ileostomy and subsequent reanastomosis | None | None | Sig | Non-infl | No | N/A | No | |
| D120 | CD | 5.6 | 21 | M | Panenteric | None | No | Yes | Multiple small bowel and colonic resections with neoterminal ileum in continuity with rectum | Vedolizumab, tofacitinib, loperamide | Pyoderma gangrenosum, chelitis, oral aphthous ulcers, fever without source | Rec | Non-infl | Yes | Opt | No | Cystic |
| D122 | CD | 5 | 7 | F | Panenteric | None | Yes | No | None | Infliximab, cholecalciferol | Arthralgias, fever without a source, oral aphthous ulcers | DC | Non-infl | No | N/A | No | |
| D125 | UC | 5.3 | 15 | F | Colonic | None | No | N/A | None | Mesalamine, cholecalciferol, lactobacillus rhamnosus | None | Sig | Non-infl | Yes | No | No | |
| D126 | UC | 2.8 | 23 | M | Pancolitis | None | No | N/A | None | Mesalamine | None | TC | Infl | Yes | No | No | |
| D127 | UC | 4 | 21 | M | Pancolitis | None | No | N/A | None | Vedolizumab, tacrolimus, balsalazide, sulfamethoxazole-trimethoprim, indigo carmine, cholecalciferol | None | Rec | Non-infl | No | N/A | No | |
| D128 | UC | 5 | 5 | M | Pancolitis | None | No | N/A | None | Sulfasalazine, methylprednisone, cortenemas | None | TC | Infl | Yes | Unk | No | |
| D130 | UC | 2.8 | 12 | F | Pancolitis | None | No | N/A | None | Mesalamine, Vancocin | None | Rec | Non-infl | No | N/A | No | |
| D131 | CD | 1.6 | 12 | F | Ileocolonic | None | No | No | None | Sulfasalazine, Mesalamine, Entyvio, bifidobacterium-lactobacillus, folic acid, vitamin D3 | None | Sig | Infl | No | N/A | No | |
| D132 | UC | 5.8 | 5.8 | F | Pancolitis | None | No | N/A | None | None | None | Sig | Infl | Yes | No | No | |
| D133 | UC | 1 | 9 | F | Panenteric | None | No | N/A | None | Adalimumab, prednisone | None | Sig | Non-infl | Yes | No | No | |
| D134 | CD | 0.5 | 2 | M | Colonic | PLCG2 | Yes | No | None | Sulfasalazine, folic acid, omeprazole, lidocaine-prilocaine topical | Erythema nodosum, pyoderma gangrenosum | Rec | Non-infl | No | N/A | No | |
| D135 | UC | 3 | 3 | M | Colonic | None | No | N/A | None | None | Fever without source | DC | Infl | No | N/A | No | |
| D139 | CD | 3 | 8 | M | Panenteric | None | No | Yes | Unk | Steroids, infliximab, adalimumab, vedolizumab | Poor weight gain, anemia, recurrent lymph node abscesses, balanitis/scarred phimosis, perioral eczema/impetigo | Sig | Infl | No | N/A | No | |

#### Supplementary Table 1. Continued

| | | | | | | | | | | | | | | | | | |
|---|---|---|---|---|---|---|---|---|---|---|---|---|---|---|---|---|---|
| D140 | IBDU | 5.9 | 18 | F | Ileocolonic | None | Yes | No | Caecal resection ileocolonic anastomosis | Azathioprine, methotrexate, infliximab, adalimumab, golimumab, ustekinumab, | Growth failure, nausea, vomiting, thalassemia | DC | Non-infl | No | N/A | No | |
| D141 | IBDU | 1 | 1.5 | F | Colonic | None | Yes | No | Unk | Intravenous immunoglobulin, various antibiotics since births, anakinra | Growth failure, constant fungal and bacterial infections, skin erosion | DC | Non-infl | No | N/A | No | |
| D142 | CD | 6 | 15 | F | Colonic | DKC1 | No | Yes | Sigma resection | Metronidazole, azathioprine, methotrexate, vedolizumab | Growth failure, arthralgia/arthritis | DC | Non-infl | No | N/A | No | |
| D143 | UC | 3 | 4.5 | M | Colonic | None | No | N/A | Unk | Exclusive enteral nutrition (Modulen IBD), tazobactam, piperacillin | Growth failure, gastroenteritis (rotavirus) | DC | Infl | No | N/A | No | |
| D145 | CD | 5 | 15 | M | Panenteric | None | No | No | Unk | Adalimumab, prednisolon, colecalciferol, vedolizumab | Fever, viral infections, hepatopathy, enuresis, pityriasis alba, herpes simplex positive | AC | Infl | Yes | Unk | No | |
| D146 | IBDU | 0.3 | 1.9 | F | Panenteric | POLA1 | No | No | Unk | Mesalazine, steroids, azathioprine | Growth failure, microcephaly | DC | Infl | No | N/A | No | |
| D147 | IBDU | 0 | .16 | M | Colonic | None | No | No | Unk | Unk | Unk | DC | Infl | No | N/A | No | |
| D148 | CD | 11 | 18 | M | Ileocolonic | None | Yes | Yes | Sigma resection | Ustekinumab, azathioprine, infliximab, adalimumab, vedolizumab | Nausea, fatigue, arthralgia | DC | Infl | Yes | Unk | No | |
| D149 | IBDU | 4 | 5 | M | Ileocolonic | None | No | Yes | Unk | Mesalazin, cortisone, Betnesol, infliximab, ciprofloxacin, metronidazol, amoxicillin | Neonatal sepsis | DC | Infl | No | N/A | No | |
| D150 | CD | .75 | 2.5 | M | Ileocolonic | None | Yes | Yes | Unk | Alfamino (EEN), 5-ASA, steroids, azathioprine, infliximab, vedolizumab | Molluscum contagiosum | DC | Infl | No | N/A | No | |
| D151 | CD | 0.6 | 0.8 | M | Ileocolonic | None | No | No | Unk | Unk | Poor weight gain | DC | Infl | Yes | Unk | No | |
| D165 | IBDU | 4 | 13 | M | Panenteric | TTC7A (Biallelic c.518G>T; p.G173V and c.1355T>C; p.L452P) | No | No | None | Immunosuppressant (Neoral), corticosteroid, omeprazole, granisetron, cidofovir, voriconazole, immunoglobulin therapy (Nanogam), amlodipine, tramadol and paracetamol | Hepatic abnormalities | Duo | Unk | Yes | Opt | Yes (N=2) | Cystic |

**Dx**, diagnosis; **SC**, Sample Collection; **P. Inv**, Perianal Involvement; **Inflamm**, Inflammation; **Org Gen**, Organoid Generation; **Org Exp**, Organoid Expansion; **M**, Male; **Unk**, Unknown; **Ile**, Ileum; **Opt**, Optimal; **F**, Female; **Rec**, Rectum; **Infl**, Inflamed; **Col**, Colon; **Sub-opt**, Sub-optimal; **AC**, Ascending Colon; **Non-infl**, Non-inflamed; **TC**, Transverse Colon; **DC**, Descending Colon; **N/A**, Not Applicable; **Cae**, Caecum; **Duo**, Duodenum; **Sig**, Sigmoid. [a] Mutations in known genes associated with IBD. [b] N=1 per time point (0, 6 and 72 hours post-stimulation), unless stated otherwise. [c] Morphology of organoid cultures, 15 days after seeding 10,000 single cells.

**Supplementary Table 2. Characteristics of Control Donors and Derived Organoid Lines**

| Donor ID | Age at sample collection | Gender | Tissue | Inflammation status | Org. Gen | Org. Exp | RNA seq[a] |
|---|---|---|---|---|---|---|---|
| D008 | 13 | M | Ile | Non-infl | No | N/A | No |
| | | | AC | Infl | Yes | Opt | Yes |
| | | | Sig | Infl | Yes | Opt | Yes |
| D010 | 2 | M | Ile | Non-infl | Yes | Unk | No |
| | | | Cae | Non-infl | Yes | Unk | No |
| | | | Rec | Non-infl | Yes | Unk | No |
| D013 | 3 | M | Duo | Unk | No | N/A | No |
| | | | Col | Unk | Yes | Opt | Yes (N=2) |
| | | | Rec | Unk | Yes | No | No |
| D019 | 3 | F | Cae | Unk | Yes | No | No |
| D020 | 3 | M | Duo | Infl | Yes | Unk | No |
| | | | Ile | Unk | No | N/A | No |
| | | | Cae | Unk | Yes | Unk | No |
| | | | Rec | Unk | Yes | Unk | No |
| D022 | 2 | F | Ile | Unk | Yes | Unk | No |
| | | | Cae | Unk | No | N/A | No |
| | | | Rec | Unk | Yes | No | No |
| D024 | 1 | F | Duo | Non-infl | No | N/A | No |
| | | | Ile | Non-infl | No | N/A | No |
| | | | Cae | Non-infl | No | N/A | No |
| D025 | 24 | F | Ile | Non-infl | No | N/A | No |
| | | | Cae | Non-infl | No | N/A | No |
| D026 | 7 | M | Ile | Non-infl | Yes | Opt | Yes |
| | | | Cae | Non-infl | Yes | Opt | Yes |
| | | | Rec | Non-infl | Yes | No | No |
| D027 | 2 | F | Duo | Non-infl | Yes | Unk | No |
| D028 | 3 | F | Ile | Non-infl | Yes | Opt | Yes |
| | | | Rec | Non-infl | Yes | Unk | No |
| D030 | 11 | M | Rec | Non-infl | No | N/A | No |
| D031 | 2 | M | Duo | Non-infl | Yes | No | No |
| D032 | 18 | M | Sig | Non-infl | Yes | Opt | Yes (N=2) |
| D033 | 2 | M | Sig | Non-infl | Yes | Sub-opt | Yes (N=2) |
| D034 | 3 | M | Col | Non-infl | Yes | Opt | Yes (N=2) |
| D035 | 3 | F | AC | Non-infl | Yes | Opt | Yes |
| D036 | 1 | M | Cae | Non-infl | Yes | Opt | Yes |
| D038 | 2 | F | Sig | Non-infl | Yes | Unk | No |
| D042 | 4 | F | Rec | Non-infl | Yes | Sub-opt | No |
| D043 | 6 | M | Ile | Non-infl | Yes | Opt | Yes |
| | | | Col | Non-infl | Yes | Sub-opt | Yes |
| D044 | 6 | M | Col | Non-infl | Yes | Opt | Yes |
| D091 | 13 | M | DC | Non-infl | Yes | Opt | Yes |

| | | | | | | | |
|---|---|---|---|---|---|---|---|
| D094 | 11 | F | Sig | Non-infl | Yes | Sub-opt | Yes |
| D099 | 2 | F | Sig | Non-infl | No | N/A | No |
| D101 | 2 | F | Rec | Non-infl | Yes | No | No |
| D102 | 2 | M | Sig | Non-infl | Yes | Opt | No |
| D103 | 4 | M | Sig | Non-infl | No | N/A | No |
| D105 | 17 | M | Ile | Non-infl | No | N/A | No |
| D107 | 17 | M | AC | Non-infl | Yes | No | No |
| D110 | 6 | M | TC | Non-infl | Yes | Opt | No |
| D111 | 7 | M | Sig | Non-infl | No | N/A | No |
| D114 | 4 | F | Col | Non-infl | Yes | Sub-opt | No |
| D118 | 4 | F | Sig | Non-infl | No | N/A | No |
| D121 | 5 | F | Sig | Non-infl | No | N/A | No |
| D123 | 6 | M | Rec | Non-infl | No | N/A | No |
| D124 | 4 | M | Sig | Non-infl | No | N/A | No |
| D129 | 1 | M | DC | Non-infl | Yes | Unk | No |
| D136 | 6 | F | Rec | Infl | No | N/A | No |
| D137 | 0 | M | Sig | Non-infl | No | N/A | No |
| D138 | 5 | F | DC | Infl | No | N/A | No |
| D144 | 17 | M | DC | Infl | No | N/A | No |
| STE076 | 8 | F | Duo | Unk | Yes | Opt | Yes |
| STE094 | 12 | M | Ile | Unk | Yes | Sub-opt | Yes |
| STE150 | 41 | F | Ile | Unk | Yes | Opt | Yes |
| HC4 | 1 | M | Ile | Unk | Yes | Opt | Yes |

**Org Gen**, Organoid Generation; **Org Exp**, Organoid Expansion; **M**, Male; **Ile**, ileum; **Non-infl**, non-inflamed; **N/A**, Not Applicable; **AC**, Ascending Colon; **Infl**, Inflamed; **Opt**, Optimal; **Sig**, Sigmoid; **Cae**, Caecum; **Rec**, Rectum; **Duo**, Duodenum; **Unk**, Unknown; **Col**, Colon; **DC**, Descending colon; **Sub-opt**, Sub-optimal; **TC**, Transverse Colon. [a] N=1 per time point (0, 6 and 72 hours post-stimulation), unless stated otherwise

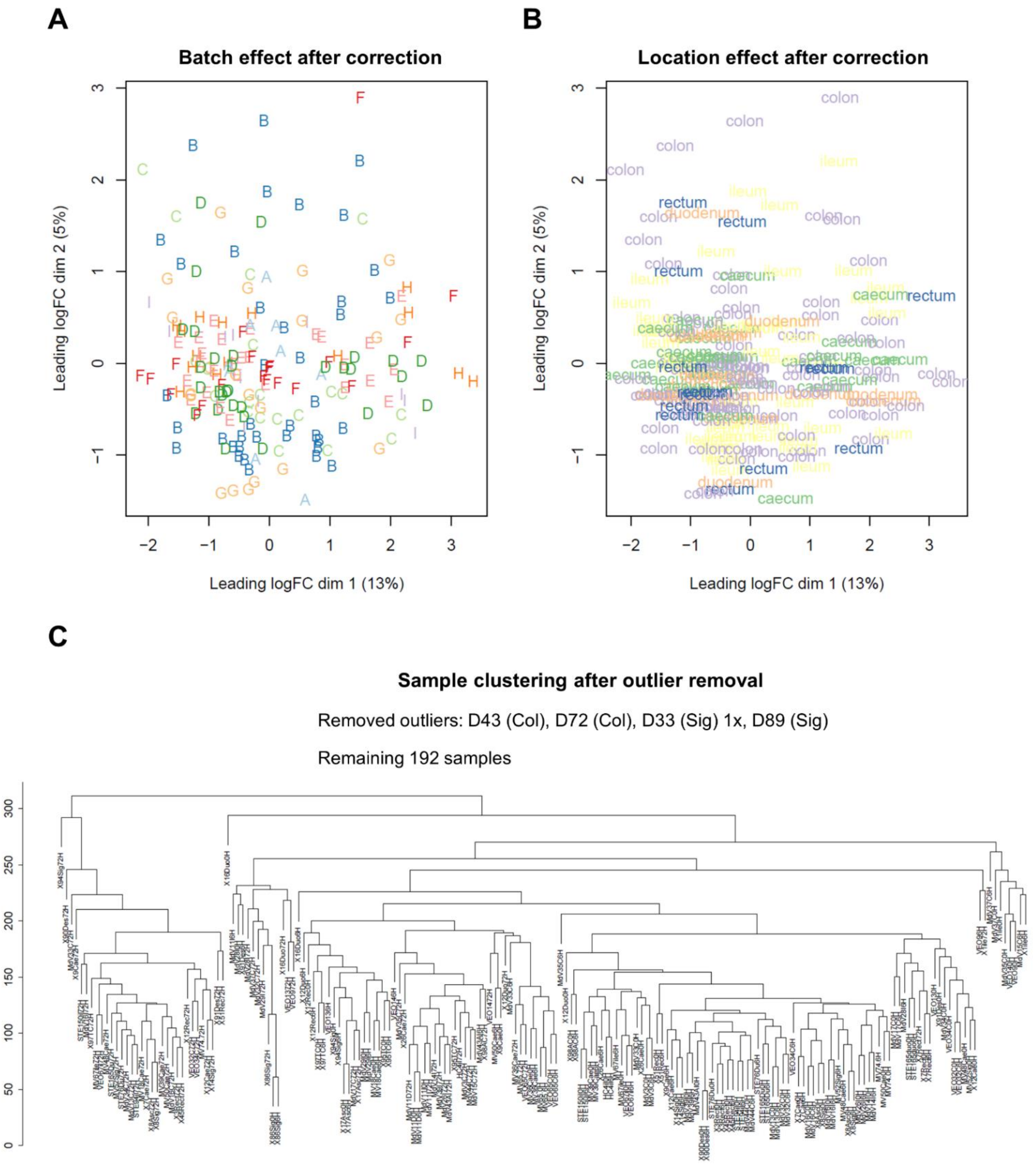


**Supplementary Figure 1. Quality control plots related to the cohort of intestinal organoid transcriptomes**. Multidimensional scaling (MDS) plots to visualize the effect of Batch **(A)** and intestinal region **(B)**. The dimensions represent the underlying factors that explain the dissimilarities among the observations with the clusters representing the groups of observations that are similar to each other. **(C)** A dendrogram showing the results of hierarchical clustering (method ='ward') of the transcriptomes of 192 remaining intestinal organoid samples after removing outlier samples: D43(col), D72 (Col), D33 (Sig) 1x, D89 (Sig)

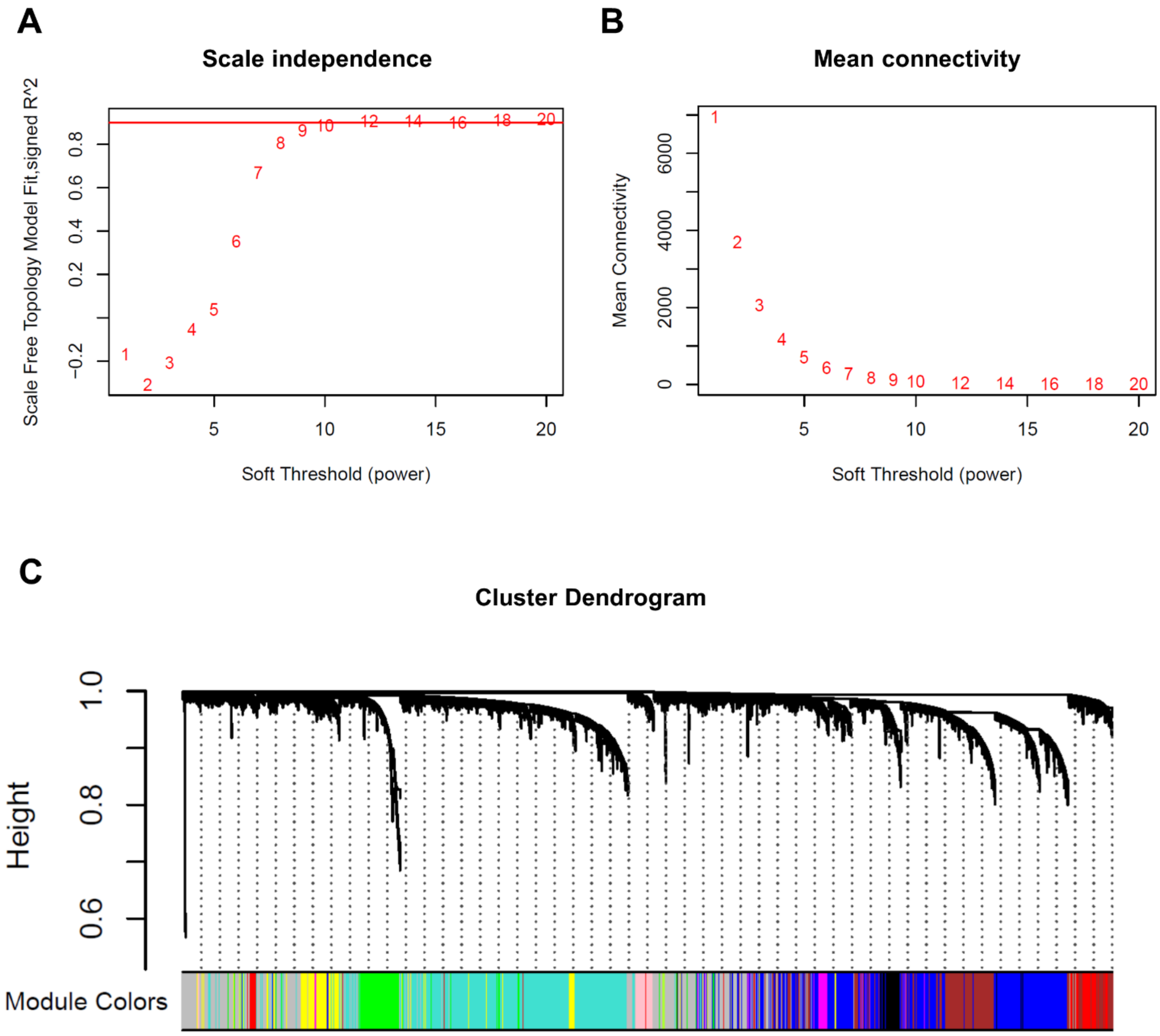


**Supplementary Figure 2. (A)** Scale independence, **(B)** mean connectivity plots and **(C)** cluster dendrogram for the signed WGCNA analysis of cohort of pediatric IBD and control intestinal organoid transcriptomes. A power threshold of 12 was chosen for the network generation. Each sample is assigned to a single module annotated by color.

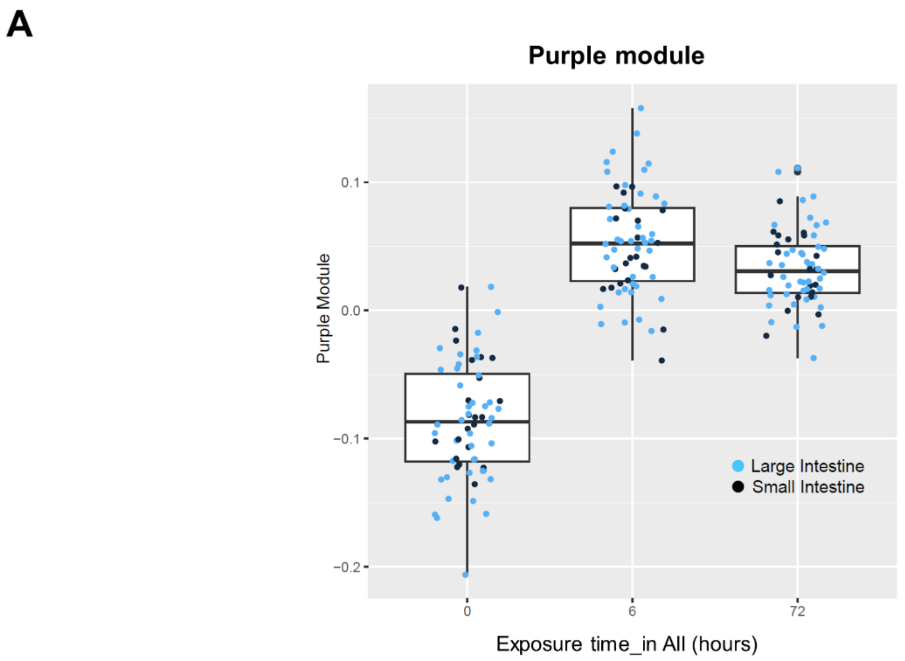


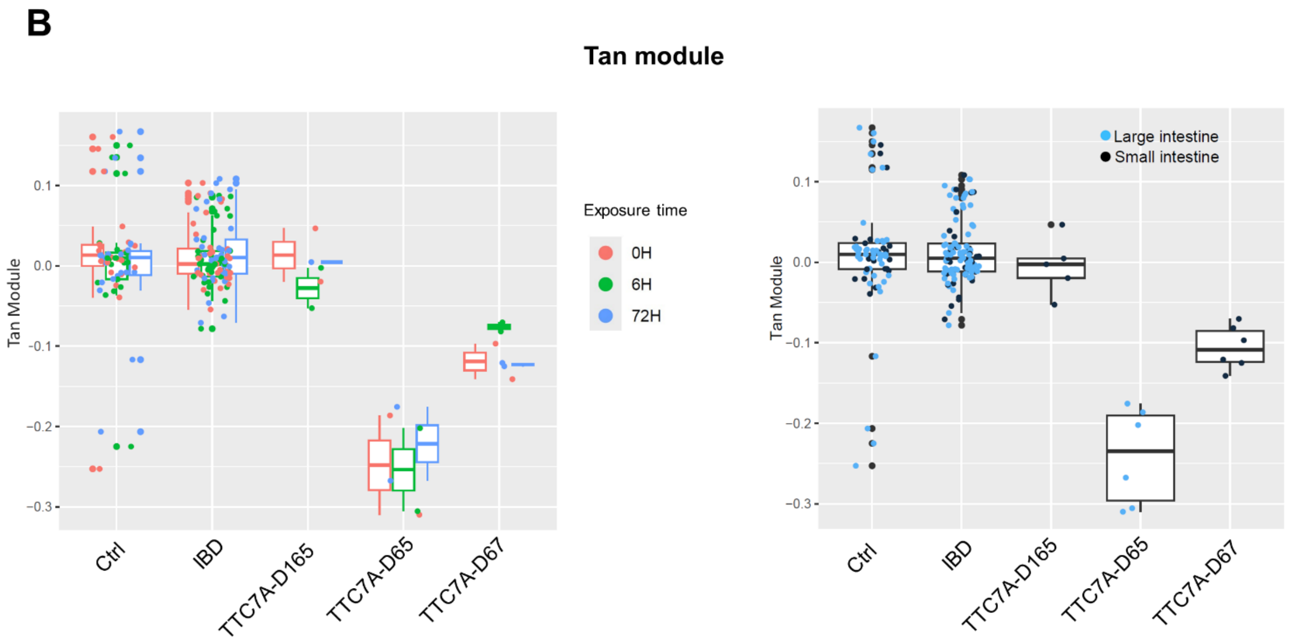


**Supplementary Figure 3. (A)** Boxplot showing the expression of the purple module at different exposure times (0, 6, and 72 hours) to bacterial lysate across approximately 60 samples from IBD and control organoid lines at each time point. Each point represents an individual sample, with color indicating intestinal location—light blue for the large intestine and dark blue for the small intestine. **(B)** Tan module expression across different conditions. The left panel shows expression levels for different conditions (Ctrl, IBD, TTC7A-D165, TTC7A-D65, and TTC7A-D67), with points colored according to exposure time (0, 6, and 72 hours). The right panel presents the same data, but points are colored by intestinal location—light blue for the large intestine and dark blue for the small intestine.
